# Modeling and treating *GRIN2A* developmental and epileptic encephalopathy in mice

**DOI:** 10.1101/737239

**Authors:** Ariadna Amador, Christopher D. Bostick, Heather Olson, Jurrian Peters, Chad R. Camp, Daniel Krizay, Wenjuan Chen, Wei Han, Weiting Tang, Ayla Kanber, Sukhan Kim, Jia Jie Teoh, Sabrina Petri, Hunki Paek, Ana Kim, Cathleen M. Lutz, Mu Yang, Scott J. Myers, Subhrajit Bhattacharya, Hongjie Yuan, David B. Goldstein, Annapurna Poduri, Michael J. Boland, Stephen F. Traynelis, Wayne N. Frankel

**Affiliations:** Columbia University Institute for Genomic Medicine; Columbia University Department of Genetics & Development; Columbia University Department of Neurology; Columbia University Department of Psychiatry; Columbia University Department of Otolaryngology and Head & Neck Surgery; The Jackson Laboratory Institute for Molecular Genetics; Emory University Department of Pharmacology; Epilepsy Genetics Program, Department of Neurology, Boston Children’s Hospital; Department of Neurology, Harvard Medical School; Department of Neurology, Children’s Hospital of Chongqing Medical University, Chongqing, 400014, China; Department of Neurology, Xiangya Hospital, Central South University, Changsha, 410013, China; Center for Functional Evaluation of Rare Variants (CFERV), Emory University School of Medicine, Atlanta, GA, 30322, USA

## Abstract

NMDA receptors (NMDAR) play crucial roles in excitatory synaptic transmission. Rare variants of *GRIN2A*, which encodes the GluN2A NMDAR subunit, are associated with several intractable neurodevelopmental disorders, including developmental and epileptic encephalopathy (DEE). A *de novo* missense variant, p.Ser644Gly (c.1930A>G), was identified in a child with DEE, and *Grin2a* knockin mice were generated to model and extend understanding of this intractable childhood disease. Homozygous and heterozygous mutant mice exhibit altered hippocampal morphology at two weeks of age, and homozygotes exhibit lethal tonic-clonic seizures in the third week. Heterozygous adult mice display a variety of distinct features, including resistance to electrically induced partial seizures, as well as hyperactivity and repetitive and reduced anxiety behaviors. Multielectrode recordings of mutant neuronal networks reveal hyperexcitability and altered bursting and synchronicity. When expressed in heterologous cells, mutant receptors exhibit enhanced NMDAR agonist potency and slow deactivation following rapid removal of glutamate, as occurs at synapses. Consistent with these observations, NMDAR-mediated synaptic currents in hippocampal slices from mutant mice show a prolonged deactivation time course. Standard antiepileptic drug monotherapy was ineffective in the patient, but combined treatment of NMDAR antagonists with antiepileptic drugs substantially reduced the seizure burden albeit without appreciable developmental improvement. Chronic treatment of homozygous mutant mouse pups with NMDAR antagonists delayed the onset of lethal seizures but did not prevent them. These studies illustrate the power of modeling severe neurodevelopmental seizure disorders using multiple experimental modalities and suggest their extended utility in identifying and evaluating new therapies.

## INTRODUCTION

N-methyl-D-aspartate receptors (NMDARs) are ligand-gated ion channels involved in brain development and fast excitatory neurotransmission (Traynelis *et al*., 2010; Paoletti *et al*., 2013). These channels are heterotetramers composed of two obligate GluN1 subunits typically paired with two GluN2 subunits that are encoded by four genes – *GRIN2A, GRIN2B, GRIN2C*, and *GRIN2D* (Traynelis *et al*., 2010; Paoletti *et al*., 2013). NMDARs can also form triheteromeric receptors through combinations of *GRIN1* and two different GluN2 subunits. Expression of these different complexes is spatially and temporally defined and varies among specific neuronal subtypes (Akazawa *et al*., 1994; Monyer *et al*., 1994; Bar-Shira *et al*., 2015). Each subunit contains a cytosolic carboxy terminal (CTD), a transmembrane region (TM), an agonist binding domain (ABD), and an extracellular amino-terminal domain (ATD) that allow modulation of receptor activity and function. NMDARs are activated by binding of glycine and glutamate to GluN1 and GluN2 respectively, leading to opening of a cation selective pore. NMDARs undergo voltage-dependent channel block by extracellular Mg^2+^, and depolarization can relieve Mg^2+^ block, allowing co-agonist activation to trigger Na^+^ and Ca^2+^ entry. The capacity of NMDARs to sense pre- and post-synaptic signals contributes to their critical role in synapse formation, development, and maintenance; they are thus key regulators of synaptic plasticity.

Pathogenic variants in *GRIN* genes are associated with developmental and early onset epileptic encephalopathies (DEE) with accompanying comorbidities, such as intellectual disability, autism, aphasia, and schizophrenia (Endele *et al*., 2010; Hamdan *et al*., 2011; de Ligt *et al*., 2012; Lesca *et al*., 2013; Petrovski and Kwan, 2013; Yuan *et al*., 2015; Hu *et al*., 2016; XiangWei *et al*., 2018). A variety of *GRIN2A* variants are associated with epilepsies, presumably due to haploinsufficiency or alteration of receptor properties. In particular, the TM domain and linker regions connecting the ABD are highly intolerant to genetic variation, as evidenced by a paucity of missense variants in these regions in control populations (Ogden *et al*., 2017). Variants that do occur in these regions, such as P552R and P557R (Ogden *et al*., 2017), L812M (Pierson *et al*., 2014), M817V (Chen *et al*., 2017), and N615K (Endele *et al*., 2010), have been reported in patients with DEE. Furthermore, whole cell and single channel patch clamp recordings from cells with heterologous expression of mutant GluN2A receptors reveal gain-of-function via enhanced agonist potency and/or prolonged channel open times (Endele *et al*., 2010; Pierson *et al*., 2014; Ogden *et al*., 2017). These findings demonstrate the critical role of the TM and linker domains in channel gating and suggest that other variants in this region could also result in altered receptor kinetics.

We describe a patient with DEE who presented with infantile spasms and impaired development in the setting of the *GRIN2A* missense variant c.1930A>G, resulting in the amino acid substitution S644G in the highly conserved third transmembrane region (TM3). We examined the variant in *Xenopus laevis* oocytes and in HEK cells, in a knock-in mouse model, and in primary neuronal networks derived from the mutant mice. S644G confers gain-of-function NMDAR characteristics. For the patient, a combination of conventional anti-seizure medications and NMDAR antagonists partially mitigated seizures but did not improve developmental comorbidities. The mutant *Grin2a*^S644G^ mice displayed several expected and unusual hyperexcitability-related features, including lethal seizures in late postnatal development. Hyperexcitability was also observed in mouse neuronal networks and was mitigated by NMDAR antagonists. In mutant mice an NMDAR combination drug delayed the onset of lethal seizures by 30%. Together with the clinical data, the parallel *in vitro*, *in vivo*, and *ex vivo* genetic model platforms provide powerful and complementary data for understanding the basis of *GRIN2A*-related disease and ultimately for evaluating potential new treatments in the broader range of patients with *GRIN2A* gain-of-function variants.

## MATERIALS AND METHODS

### Patient ascertainment and phenotyping

The patient was referred for consultation to the Epilepsy Genetics Program at Boston Children’s Hospital (BCH) after clinical genetic testing for DEE revealed the *de novo* pathogenic variant. Consent was obtained to share deidentified patient data with collaborating researchers under a protocol approved by the Institutional Review Board (IRB). After discussion with the Office of Regulatory Affairs and IRB, the patient was treated clinically with off-label memantine and dextromethorphan based on published reports of use for other indications and safety in children. The family was counselled that use of these medications was off-label. Clinical management with other medication and standard clinical monitoring with seizure counts, developmental assessment, and EEG continued as per standard of care.

### Mice, husbandry, and genotyping

All mice were bred and procedures conducted at The Jackson Laboratory, Columbia University Irving Medical Center, or Emory University, each fully accredited by AAALAC, approved by respective IACUC and performed in accordance with state and federal Animal Welfare Acts and Public Health Service policies.

*Grin2a^S644G^* mice (official symbol: *Grin2a*^em1(S644G)Frk^) were generated in the C57BL/6NJ (B6NJ) mouse strain using CRISPR/Cas9 and an oligonucleotide donor sequence as part of the JAX Center for Precision Genetics (JCPG) and maintained by backcrossing heterozygous males to wildtype B6NJ females. For some experiments, as noted, male mutants were crossed with FVB/NJ females to generate cohorts of F_1_ or F_2_ hybrid mice. Mice were housed in ventilated cages at controlled temperature (22–23°C), humidity ~60%, and 12h:12h light:dark cycles. Mice had *ad libitum* to regular chow and water. Mice were genotyped using the following PCR amplification primers (540bp amplicon), using standard thermocycler conditions and Sanger sequenced (GeneWiz, Inc); F:CGAGTGTACACGCTGTGGAAATAG; R: TCAACCAGTGCTACAGAGTGATCT.

### Seizure studies

#### Pup spontaneous seizure observation

Events in each cage were recorded live using above cage Sony HD digital cameras connected to a Samsung VCR set (Hanwha Techwin Co., LTD) for 24 hours.

#### Video-EEG

Electrode implantation of adult mice was performed surgically essentially as described (Asinof *et al*., 2016). Three silver wire electrodes were placed subdurally 1 mm rostral to Bregma and 1 mm to either side of midline, and another over the cerebellum as reference. Signal was acquired on a Quantum 128 research amplifier (Natus, Inc). Differential and referential montages were examined in Neuroworks software (Natus).

#### Electroconvulsive threshold

Tests were performed using transcorneal electrodes with the Ugo Basile Model 7801 electroconvulsive device as described previously (Asinof *et al*., 2015). Settings for minimal clonic forebrain seizures were 299 Hz, 1.6 ms pulse width, 0.2 s duration, variable current. Data were analyzed as the integrated root mean square (iRMS, area under the curve) as (square root of Hz) × current × pulse width × duration.

### Histology

Postnatal day 14 pups were perfused transcardially using Bouin’s solution. Brains were fixed, embedded in paraffin, sliced and stained with hematoxylin and eosin using standard methods.

### mRNA isolation and RT-qPCR

RNA from whole brains was extracted as previously described (Johnstone *et al*., 2010). cDNA was synthesized using a qScript cDNA synthesis kit (Quanta BioSciences) using 1μg per sample. Quantitative RT-qPCR was performed on a Quant Studio 5 (Applied Biosystems). Primers were designed using Primer3. Melt curve analyses were used to eliminate primer-dimer artifacts and check reaction specificity. All data were normalized to *Gapdh*. Primer sequences for selected targets: *Grin2b* – F: GAGCACGTGGACTTGACTGA; *Grin2b* – R: TACCACTCCGTGCTTATCGC; *Grin2a* – F: TCTTGAACTACAAGGCCGGG; *Grin2a* – R: GCGAGGTCAATCTGCCTCTT; PSD95 – F: ACCAAGATGAAGACACGCCC; PSD95 – R: TCCGTTCACCTGCAACTCAT; *Gapdh* – F: GACCACAGTCCATGCCATCAC; *Gapdh* – R: GTCCACCACCCTGTTGCTGTA

### Protein analysis

Crude membrane extracts were prepared by homogenizing cortices in 0.32 M sucrose, 1 mM NaHCO3, 1 mM MgCl_2_; and a protease inhibitor cocktail (ThermoScientific Cat#A32955). Protein concentrations were determined by Bradford’s method using BSA as standard. Samples (25 μg) were diluted in 4× sodium dodecyl sulphate (SDS) protein gel loading dye, boiled for 5 min, separated on 10% Mini-PROTEAN TGX gel (Bio-Rad) and electroblotted to PVDF transfer membrane (Millipore, Cat# ISEQ00010). Nonspecific binding was blocked for 30 mins or 1 h at room temperature with 5% nonfat dry milk in Tween-Tris-buffered saline (TBS-T). Membranes were incubated overnight at 4 °C with the following antibodies: NMDAR2A (Millipore, Cat# AB1555P), NMDAR2B (Cell Signaling, Cat# 4207S), all normalized to Tubulin β-3 (BioLegend, Cat# 802001), PSD95 (Pierce or Synaptic Systems). Secondary antibodies anti-mouse and antirabbit (Cat# SA00001-1 and SA00001-2, Proteintech) were diluted at 1:10,000. All antibodies were prepared in 5% nonfat dry milk solution in TBS-T. Blots were developed using enhanced chemiluminescence ECL technique (Azure Biosystems, Inc, Model C400) and relative band densities were quantified using ImageJ software.

### Mouse neurobehavioral tests

#### Open field exploration

Each mouse was placed gently in the center of a clear Plexiglas arena (27.31 x 27.31 x 20.32 cm Med Associates ENV-510) lit with dim light (~ 5 lux), and allowed to ambulate freely for 60 min. Infrared (IR) beams embedded along X, Y, Z axes of the arena automatically track distance moved, horizontal movement, vertical movement, stereotypies, and time spent in center zone. After testing, mice were returned to the home cage and the arena cleaned with 70% ethanol followed by water, and wiped dry.

#### Elevated plus maze

This test was conducted as described previously (Yang *et al*., 2012). The maze consists of two open arms (30cm x 5cm) and two closed arms (30 x 5 x 15 cm) extending from a central area (5 x 5 cm). Photo beams embedded at arm entrances register movements. Room illumination was approximately 30 lux. Mice were placed in the center, facing a closed arm and allowed to freely explore the maze for 5 min. Time spent in open and closed arms, the junction, and number of entries into each, were scored automatically by MED-PC V 64bit Software (Med Associates). After testing, the mouse was gently removed from the maze and returned to home cage. The maze was cleaned with 70% ethanol and wiped dry between subjects.

#### Repetitive behaviors

Each subject was placed in an empty clean mouse cage, with a 0.25 cm layer of clean bedding lining the bottom. The cage was covered with a metal wire lid and a plastic filter top. Behaviors were recorded for 60 minutes and analyzed using the time-sampling method for the last 10 minutes. An observation was scored every 30 seconds for a total of 20 samples. Occurrences of circling, hanging on the wire, backflipping, self-grooming, resting, and exploration were recorded for each sample.

#### Social interaction

The test was conducted in the PhenoTyper chamber (Noldus) as previously described (Yang et al., 2012). The floor of the arena was covered with a 0.5 cm layer of clean bedding. Each subject mouse was singly housed in a clean cage for one hour before the test. After this brief isolation, the freely moving subject mouse and freely moving age- and sex-matched B6 partner mouse were placed simultaneously in the arena and interactions were videotaped for 10 min. Social interactions were scored by a highly trained observer blind to genotype. Time spent in nose-to-nose sniff, front approach, follow, nose-to-anogenital sniff, and push-crawl were scored and summed to yield % time social interaction.

#### Acoustic startle response

Acoustic startle response was tested using the SR-Laboratory System (San Diego Instruments, San Diego, CA) as described previously (Yang et al., 2012). Test sessions began by placing the mouse in the Plexiglas holding cylinder for a 5-min acclimation period. For the next 8 min, mice were presented with each of 6 trial types across 6 discrete blocks of trials, for a total of 36 trials. The intertrial interval was 10–20 s. One trial type measured the response to no stimulus (baseline movement). The others measured *startle* response to 40 ms sound bursts of 80, 90, 100, 110 or 120 dB. The six trial types were presented in pseudorandom order such that each was presented once within a block of six trials. Startle amplitude was measured every 1 ms over a 65 ms period beginning at the onset of the startle stimulus. The maximum startle amplitude over this sampling period was taken as the dependent variable. Background noise level of 70 dB was maintained over the duration of the test session.

#### Auditory Brainstem Response (ABR)

Mice were anesthetized with ketamine (100 mg/kg, i.p.) and xylazine (10 mg/kg, i.p.). ABR clicks test was performed in an acoustically and electrically shielded booth (Industrial Acoustic Company, Bronx, NY, USA). The sound intensity level decreased from 80 to 5dB at 5 dB intervals. The electrodes were placed subcutaneously on the vertex (active), mastoid (reference), and hindlimb (ground). The threshold was determined as the lowest intensity level that induced a detectable ABR response. Measurements were recorded and analyzed with a S3 ABR system (Tucker Davis Technologies, Alachua, FL, USA).

### Multielectrode array (MEA)

#### Primary neuron culture

Prior to use 48-well MEA plates (Axion BioSystems) were coated overnight with 50 μg/mL poly-D-lysine (Sigma) in 0.1M borate buffer (pH 8.5). Cortices from littermates were incubated in activated 20 U/mL Papain/DNase (Worthington) for 15 minutes at 37°C, centrifuged at 300 x *g* for 5 min, and washed in 1X PBS. Cell pellets were suspended in NBA/B27 [consisting of Neurobasal-A (Life Technologies), 1X B27 (Life Technologies), 1X GlutaMax (Life Technologies), 1% HEPES, and 1X Penicillin/Streptomycin], supplemented with 1% fetal bovine serum (Gibco) and 5 μg/mL laminin. Fifty thousand cells per well were plated on pre-coated MEA plates in a 40 μl convex drop yielding a density of ~1600 cells/mm^2^. The day after plating media was removed and replaced with pre-warmed NBA/B27. Cultures were maintained at 37°C in 5% CO_2_. Medium was 50% changed every other day with NBA/B27 starting on DIV3.

#### MEA data collection and analysis

Recordings were conducted prior to media change for 15 minutes per day using Axion BioSystems Maestro 768 channel amplifier and Axion Integrated Studios software (v2.4) at 37°C in a carbogen mixture of 95% O_2_, 5% CO_2_. A Butterworth bandpass filter (200-3000 Hz) and an adaptive threshold spike detector set at 7x the standard deviation of noise to record raw data and spike list files. Data were analyzed using the meaRtools package (Gelfman *et al*., 2018). Spike list files were used to extract additional spike, burst, and network features. At least 5 spikes/minute per electrode were required; wells with fewer than 4 active electrodes for more than 30% of the total recording days were discarded, resulting in 94% of wells and 94% of electrodes over seven 48-well MEA plates deemed active.

#### Burst detection

The Maximum Interval burst detection algorithm (Neuroexplorer software, Nex Technologies) was used. Based on previous studies (Johnstone et al. 2010; Mack et al. 2014), we required that a burst consist of at least 5 spikes, lasting at least 50 ms, and that the maximum duration between two spikes within a burst was 0.05 s. Adjacent bursts were merged if the duration between them was < 0.11 s.

#### Network burst detection

Spike time within spike trains from all electrodes in a well was normalized using a bin size of 2 ms to guarantee that at most one spike is called within each bin. A Gaussian filter was applied to smooth the binned spike train from each electrode. The smoothed signal was then standardized to a maximum value of 1. The sum of all smoothed signals from individual electrodes were then smoothed again using the same Gaussian filter before applying the Otsu global thresholding method (Otso, 1975) to the signal from each well to automatically detect burst intervals. Network bursts were analyzed in 400 ms windows. Mutual information was calculated as previously described (Gelfman *et al*., 2018).

#### Electrical stimulation

After DIV25 two electrodes per well (370 um apart) were probed with ten biphasic 0.2 Hz pulses sequentially to investigate evoked electrical responses. Evoked responses were measured with a pulse amplitude fixed at 1.6-2.0 V peak-to-peak with the positive phase coming before the negative phase, and lasting for 200 μs per phase (Wagenaar *et al*., 2004; Wagenaar and Potter, 2004). Peristimulus time histograms (PSTH) were generated using the Axion BioSystems Neural Metric Tool (v2.2.4). A PSTH for each well was calculated by recording the spiking activity over a 400 ms period after each stimulation (20 stimuli per well) then averaging the number of spikes on active electrodes (>0.1 Hz) in the well per trial for each histogram bin (1 ms) (Shahaf and Marom, 2001).

### Two-electrode voltage clamp (TEVC) and whole-cell voltage clamp recordings

cDNA for wildtype human NMDA receptor subunits GluN1-1a (hereafter GluN1 (NCBI NM_007327/ NP_015566) and GluN2A (NCBI NM_000833/NP_000824) were subcloned into pCI-neo (Promega, Madison, WI). Site-directed mutagenesis was performed to generate human GluN2A-S644G construct as previously described (Yuan *et al*., 2015). Preparation and injections of cRNA, TEVC from *Xenopus laevis* oocytes, and expression of triheteromeric receptors were performed as previously described (Hansen *et al*., 2014). cDNAs encoding wild type and mutant GluN2A subunits were fused at the C-terminal with a synthetic linker, a coiled-coil domains C1 and C2, and an ER retention signal as previously described (Hanson et al., 2014). cRNAs encoding GluN1, as well as C1- and C2-tagged GluN2A, were injected at a 1:2:2 ratio with ~10 ng total cRNA at a total volume of 50 nl (1:2:2 for GluN1:GluN2A_c1_:GluN2A_c2_, or GluN1:GluN2A_c1_-S644G:G1uN2A_c2_ or GluN1:GluN2A_c1_-S644G:GluN2A_c2_-S644G) and were kept at 19°C in Barth’s solution for 1 to 3 days before recording. For all experiments, the cRNA was diluted with RNase-free water. Barth’s solution contained (in mM) 88 NaCl, 2.4 NaHCO_3_, 1.0 KCl, 0.33 Ca(NO_3_)_2_, 0.41 CaCl_2_, 0.82 MgSO_4_, and 5 Tris/HCl (pH 7.4). Glutamate and glycine concentration-response curves were recorded at −40 mV holding potential. Magnesium and drug inhibition curves were recorded at −60 mV, and zinc inhibition curves were recorded at −20 mV (pH 7.3, Yuan et al., 2014). The oocyte recording solution contains (in mM) 90 NaCl, 1 KCl, 10 HEPES, 0.5 BaCl_2_, and 0.01 EDTA (pH 7.4). HEK293 cells were transfected with GFP, GluN1, and GluN2A-wildtype, GluN2A-S644G, GluN2A_c1_, GluN2A_c2_, GluN2A_c1_-S644G, and/or GluN2A_c2_-S644G using the calcium phosphate method (Chen and Okayama, 1987). Whole-cell voltage clamp were performed with a rapid solution exchange system and analyzed by ChannelLab.

### Hippocampal whole-cell patch clamp recordings

Horizontal hippocampal brain slices (300 *μ*m) were made from postnatal day 13-17 wildtype (+/+) and S644G/+ littermate F_2_ hybrid mice of both sexes using a vibratome (TPI, St. Louis, MO). During preparation, the slices were bathed in an ice-cold (0-2°C) sucrose-based artificial cerebral spinal fluid aCSF containing (in mM) 230 sucrose, 24 NaHCO_3_, 10 glucose, 2.5 KCl, 1.25 NH_2_PO_4_, 10 MgCl_2_, and 0.5 CaCl_2_ bubbled with 95% O_2_/5% CO_2_. After preparation, slices were transferred to a NaCl-based recovery aCSF containing (in mM) 130 NaCl, 24 NaHCO_3_, 10 glucose, 2.5 KCl, 1.25 NH_2_PO_4_, 4 MgCl_2_, and 0.5 CaCl_2_ bubbled with 95% O_2_/5% CO_2_. Recovery solution was supplemented with 30 μM 7-chlorokynurenic and incubated at 32°C for 30 minutes before returning them to room temperature and left to recover for at least 1 hour prior to use.

Evoked CA1 pyramidal cell EPSC were recorded at room temperature in aCSF but with 0.2 mM MgCl_2_, 1.5 mM CaCl_2_, and no 7-chlorokynurenic acid. Thin-walled borosilicate glass (1.5-mm outer, 1.12-mm inner diameter, WPI Inc., Sarasota, FL) was used to fabricate recording electrodes (4–6 MΩ), which were filled with (in mM) 100 Cs-gluconate, 5 CsCl, 0.6 EGTA, 5 BAPTA, 5 MgCl_2_, 8 NaCl, 2 Na-ATP, 0.3 Na-GTP, 40 HEPES, 5 Na-phosphocreatine, and 3 QX-314. Membrane resistance was on average 239+39 (+/+) and 219+36 MΩ (+/S644G), series resistance was on average 15+1.1 (+/+) and 17+1.6 MΩ (+/S644G), and cell capacitance was 110+5.9 (+/+) and 120+9.3 pF (+/S644G). Monosynaptic release of glutamate was evoked at 0.033 Hz using a monopolar tungsten-stimulating electrode by injecting 50–150 μA of current for 0.1 ms along the Schaffer collateral pathway. CA1 pyramidal cells were held at −30 mV to minimize magnesium block. The NMDAR-mediated current component was isolated via bath application of 10 μM NBQX (AMPAR antagonist, Sigma) and 10 μM gabazine (GABAAR antagonist, Sigma). Cells were held in 10 μM NBQX and 10 μM gabazine for 10 minutes before baseline current responses were recorded. A total of eight epochs were recorded and averaged together. At the conclusion of recording, 200 μM DL-APV (Sigma) was applied to ensure responses were mediated via NMDARs; DL-APV caused a similar change in the baseline leak current of 15+3.2 (+/+) and – 22+7.7 pA (+/S644G).

Recordings were made using an Axopatch 1D amplifier (Molecular Devices), digitized at 20 kHz using a Digidata 1440a and Axon pClamp10 software, and filtered at 5 kHz using an eight-pole Bessel filter (−3 dB; Frequency Devices). Series resistance was monitored throughout the experiment and was typically 8–20 MΩ. If the series resistance changed >20% during the experiment, then the cell was excluded. All data were analyzed offline using ChanneLab (Synaptosoft) and synaptic decay was calculated using a dual-component exponential.

### Pharmacology

Dimethyl sulfoxide (DMSO) was from Fisher Scientific, NMDA was from Tocris, radiprodil was from Cayman Chemical, memantine and dextromethorphan were from Sigma-Aldrich. For use in MEA s tock solutions of 1000X and the final desired concentration was prepared for each compound in deionized water (NMDA 100 mM, Dextromethorphan and memantine 50 mM). MEA plates were dosed by addition of 5 uL of appropriately diluted stock concentration of drug to a parallel dosing plate, then half (150 uL) of the media from the MEA plate was removed and added to the parallel dosing plate and mixed prior to adding the media back to the MEA plate. Plates were allowed to equilibrate briefly (2-3 minutes) post administration of drug prior to recording and non-treated wells were used to verify stabilization was achieved. Concentration-response relationships were determined in a cumulative manner, in which the concentration of drug present in the medium was increased in a stepwise manner (Novellino et al 2011).

For use in mice, dextromethorphan was administered in a 0.9 % saline vehicle (at 25 mg kg^-1^) and solubilized. Formulation to dissolve radiprodil consisted of 2% N,N-dimethylacetamide (DMA, Sigma Cat# D5511)/10%propylene glycol (PG, Sigma Cat#P4347)/30% 2-hydroxypropyl-beta-cyclodextrin (HPBCD, Sigma Cat# 332593). Quinidine (Sigma Aldrich) was diluted in saline solution and administered at 10 mg/kg. Dissolved radiprodil was diluted in saline solution and administered at 2 mg kg^-1^. Nuedexta was prepared as a formulation of 25:10 mg kg^-1^ dextromethorphan: quinidine.

#### Statistics

For animal studies, the nonparametric Mann-Whitney rank-sum test was performed when data were not normally distributed or when fewer than 10 animals were available per group and another test was not more appropriate. Two-way repeated measures ANOVA was used for open field arena tests. Survival curves were analyzed using the Mantel-Cox log-rank test. Student’s |t|-test was used for hippocampal morphometry and for *in vitro* electrophysiology. For MEA neuron culture spontaneous phenotypes, data were rank- and normal quantile transformed, a least squares regression model performed, *p*-values for each DIV were obtained for plate ID and genotype effects and the latter were adjusted by Bonferroni correction for the 11 DIV measurements per feature and for DIVs with significant genotype effects, the significance between each homozygous or heterozygous against wildtype was determined by a post-hoc Dunnet’s test set at p<0.05. For MEA drug responses, data were transformed as above and genotype x treatment interaction effects were tested in a least-squares regression model. All studies were designed so that an effect size of at least 1.6 was detected at 80% or greater power.

## RESULTS

### A *de novo GRIN2A* variant in a child with DEE

#### Clinical presentation and etiology

A 4-month-old boy presented with concerns for visual tracking and developmental delay and developed seizures at 6-8 months. There were no risk factors for epilepsy nor family history of epilepsy. Prenatal history was notable for polyhydramnios and preterm labor at 32 weeks’ gestation, and birth was at 36 weeks’ gestation by Caesarean section. He required minimal resuscitation with Apgar scores of 3, 5, and 8 at 1, 5 and 10 minutes, respectively, and was monitored in neonatal intensive care for two days for feeding and swallowing concerns. MRI at 7 months was normal.

At 6-8 months the boy exhibited clusters of myoclonic seizures that evolved to infantile spasms. Other seizure types eventually included focal motor (tonic) seizures, focal impaired awareness seizures, generalized tonic seizures, and generalized tonic-clonic seizures. Electroencephalogram (EEG) recordings were consistent with epileptic encephalopathy, including diffuse background slowing and disorganization, electroclinical seizures, some with generalized and some with focal onset, and interictal generalized spike-slow wave complexes and multifocal spikes. He had profound global developmental delay without regression, cortical visual impairment, spastic quadriparesis, and oropharyngeal dysphagia warranting exclusive gastrostomy feeding. At 6 years of age, he continues to be nonverbal and non-ambulatory with limited purposeful hand use and lack of independent head control. Together, his features constitute a DEE.

Molecular diagnosis via exome sequencing identified the *de novo* heterozygous missense variant c.1930A>G in the *GRIN2A* gene encoding the GluN2A NMDAR subunit. The resulting *GRIN2A* S644G amino acid substitution resides in the transmembrane domain TM3, an area with high sequence conservation across all NMDAR genes in human and mouse (**Fig. 1a**).

**Figure 1.**
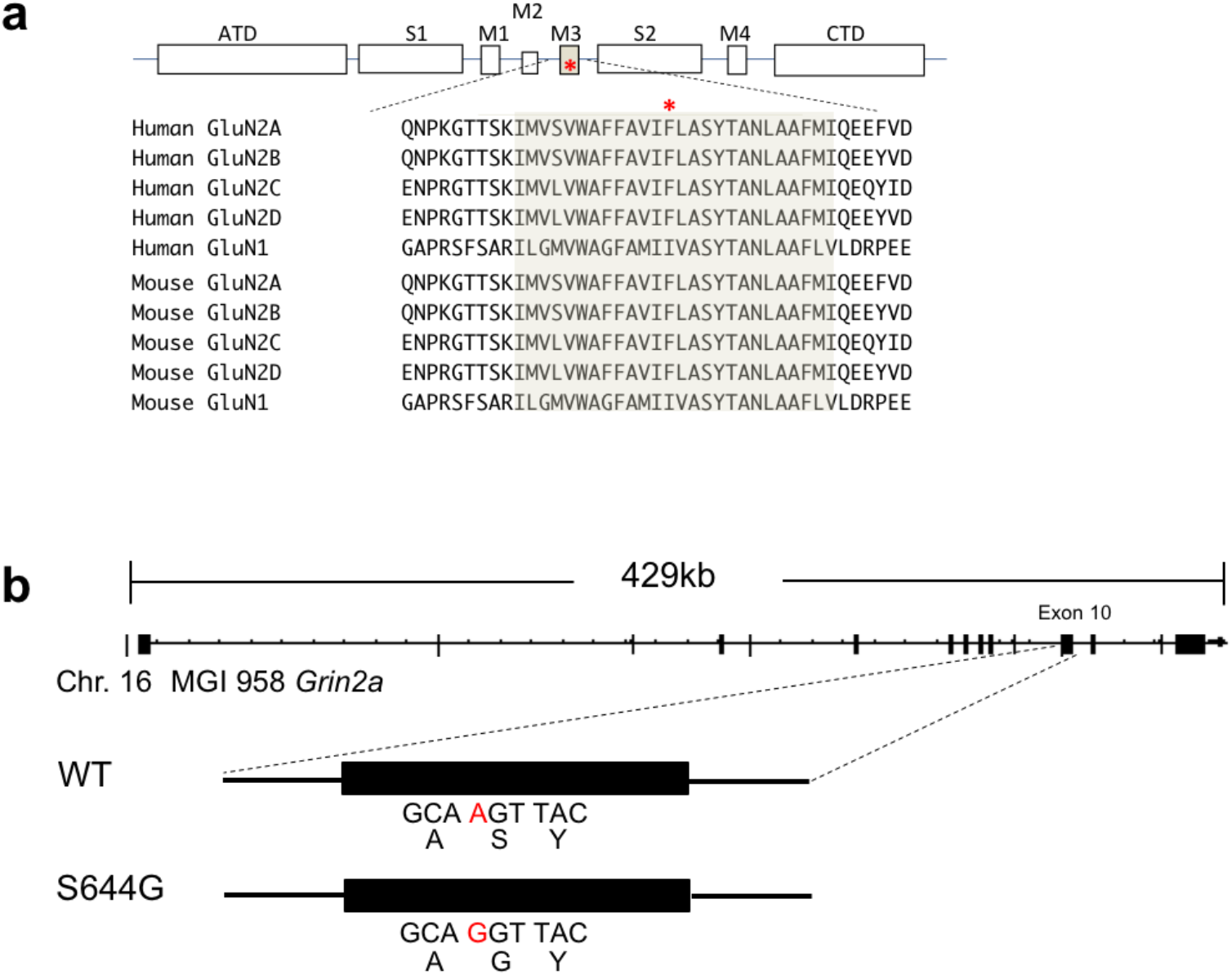
GRIN2A/Grin2a S644G substitution in human and in mouse model. (a) Alignment of the GluN2A protein sequence across human and mouse GluN subunits. The TM3 transmembrane domain is highly conserved within the NMDAR family, including the serine residue that appears mutated in the patient. ATD, amino terminal domain; S1 and S2, polypeptide chains that form the agonist-binding domain (ABD); M1, M2, M3, and M4, transmembrane domain (TM) helices 1, 3, and 4, and the membrane re-entrant loop 2; CTD, carboxy-terminal domain (COOH). (b) Structure of the mouse *Grin2a* gene and A>G nucleotide mutation that results in serine to glycine amino acid substitution induced by CRISPR/Cas9 mutagenesis creating the S644G allele from wildtype.

### *Grin2a*^S644G^ mutant mice have unusual seizure, behavior, and histopathological features

We generated a mouse line carrying the orthologous mutation on the B6NJ strain background (**Fig. 1b**). B6NJ Grin2a^S644G/+^ (S644G/+) females are fertile and had no difficulties delivering, but did not care for the pups beyond the first postnatal day (PND), regardless of pup genotype (**Supplementary Fig. 1**). Observational assessments including pup retrieval, self-grooming and pup care suggested impaired maternal behaviors in B6NJ S644G/+ females.

To study homozygous mutants, F_2_ hybrid mice were generated from an intercross between (FVB/NJ × B6NJ-S644G)F_1_ parents. Although all *Grin2a* genotypes survived to the third week (**Supplementary Fig. 1**), all S644G homozygotes unexpectedly died of a lethal seizure between PND15-17. Evidence for seizure included maximal hind limb extension, observed both in carcasses and in animals prior to death (**Fig. 2a**). Seizures began suddenly with a wild run followed rapidly by a tonic-clonic phase and ultimately maximal tonic hind limb extension, which was terminal. The early third week onset correlates with the known temporal peak of *Grin2a* expression (Bar-Shira *et al*., 2015). Heterozygous animals had a normal lifespan and neither exhibited behavioral seizures during routine handling, nor epileptiform activity during continuous video-EEG analysis (**Supplementary Fig. 2**).

**Figure 2.**
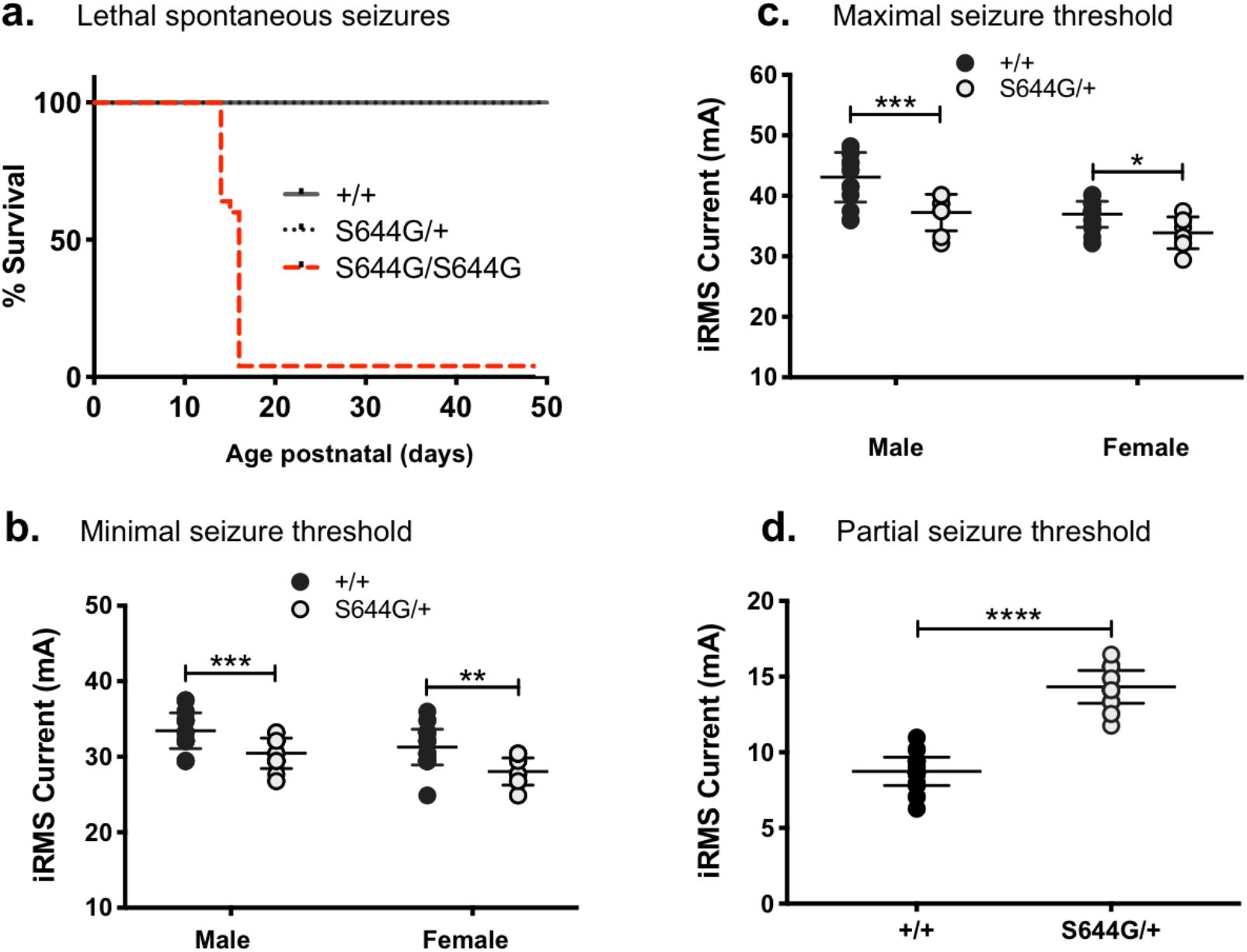
Seizure susceptibility of *Grin2a*^S644G^ mice. (a) Survival curve showing the rate of genotype-dependent death in F_2_ mice. Both male and female mice homozygous for S644G have spontaneous lethal seizures at P15-P17. (b, c) Heterozygous adult mice have lower seizure threshold in minimal seizure endpoints (female N=20 wt, 22 het, *p*=0.0048; male N=31 wt, 27 het, *p*=0.0035); and maximal seizure ECT endpoints (female N=6 het, 7 wt; *p*=0.0434; male N=15 het, 9 wt; *p*=0.0007), but (d) are significantly more resistant to partial seizures in the 6 Hz ECT test (N=33 het, 31 wt;*p*=4.1×10^-12^); *p<0.05; **p<0.01; ***p<0.001; ****p<0.0001, Mann-Whitney rank-sum test. Error bars indicate S.E.M.

Seizure susceptibility of S644G/+ adults was assessed using electroconvulsive threshold (ECT) testing targeting minimal clonic (forebrain), maximal tonic (hindbrain/brainstem), and psychomotor seizure (6 Hz test) endpoints. Heterozygous S644G/+ mice exhibited a decreased seizure threshold to both minimal and maximal seizures, (**Fig. 2b-c**), but in the 6 Hz seizure test showed an unusual resistance, suggesting significant abnormality in the limbic system (**Fig. 2d**).

*Grin2a* mRNA and protein were present but modestly decreased in heterozygous and homozygous pups at PND14, as was the scaffold PSD95 protein but not its transcript (**Supplementary Fig. 3**). *Grin2a* is expressed highly in the hippocampus and we detected thinning of the dentate gyrus and CA1 region as a common feature of S644G heterozygotes and homozygotes compared to age-matched wildtype (**Fig. 3**). We did not detect gross changes in cortical lamination. Since *Grin2a* expression is predominantly postnatal, together these results suggest the likelihood of hippocampal cellular atrophy prior to PND14.

**Figure 3.**
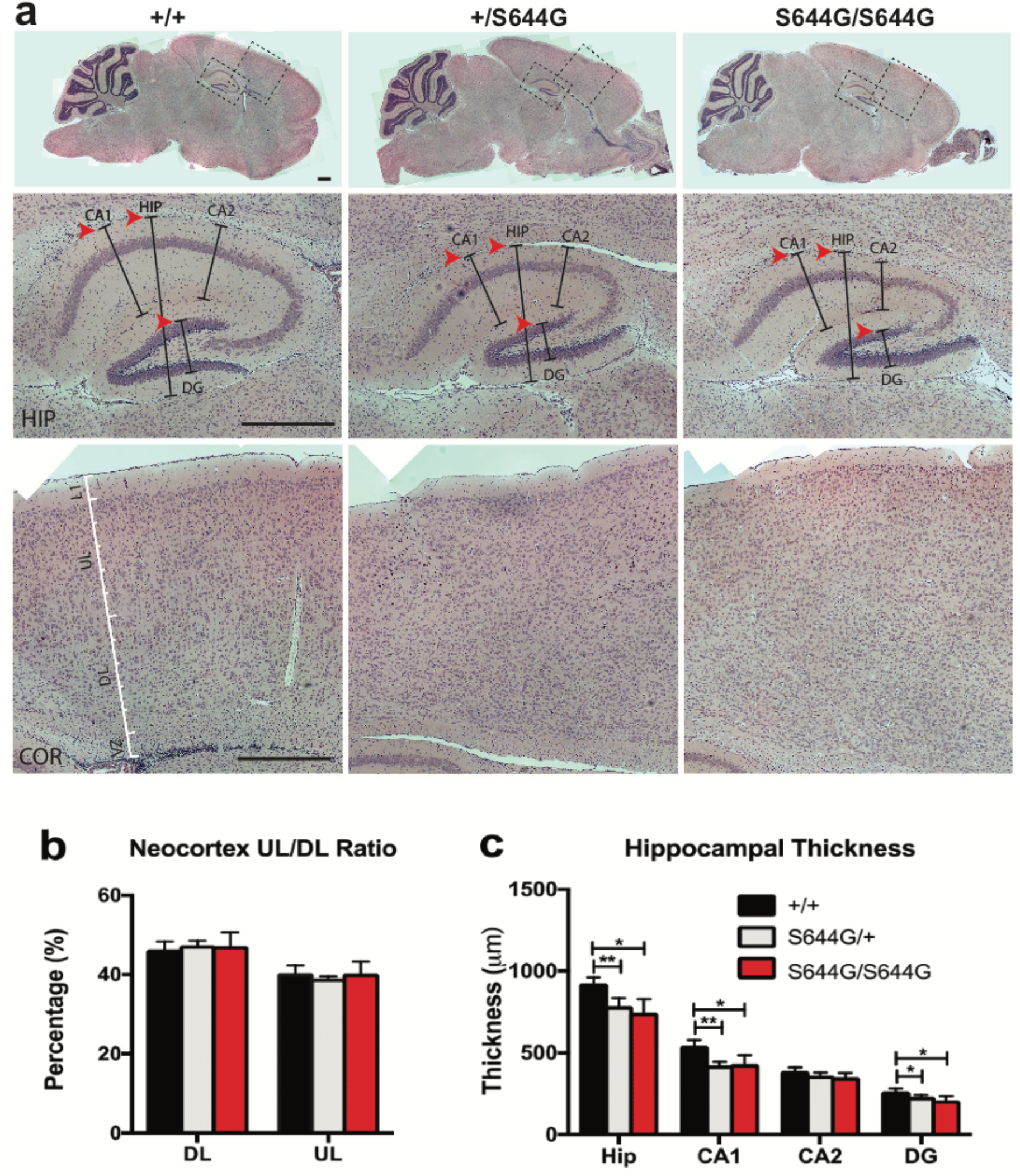
Aberrant hippocampal morphology of *Grin2a*^S644G^ adolescent mice. (a) H&E stained brain sections (Scale bars 500 μm). At PND14, markedly decreased hippocampal thickness, particularly in CA1 and DG (arrowheads). (b, c) Plot of structure thickness, showing no differences in the neocortex, but decreased thickness in the hippocampus. *p<0.05; **p<0.01; Student’s latest.

We performed a battery of neurobehavioral tests in adult littermates on both B6NJ and F_1_ hybrid backgrounds. In the open field, B6NJ S644G/+ heterozygotes of both sexes showed increased ambulation compared to wildtype (**Fig. 4a**), but in F_1_ hybrids only females displayed this phenotype (**Supplementary Fig. 4a**). B6NJ S644G/+ females also spent more time in the center of the arena, with males trending toward the same pattern (**Fig. 4b**), suggestive of decreased anxiety-like behaviors. Again, in F_1_ hybrids only females showed significantly more time in the center (**Supplementary Fig. 4b**). Genotype effects on vertical activity were not significantly different (**Supplementary Fig. 4c-d**). In the elevated plus maze, while S644G/+ mice showed no difference in total arm entries or time at the junction, indicating normal locomotor activity, S644G/+ mice of both backgrounds trended towards more time on the open arms, suggestive of decreased anxiety-like behavior (**Supplementary Fig. 5**).

**Figure 4.**
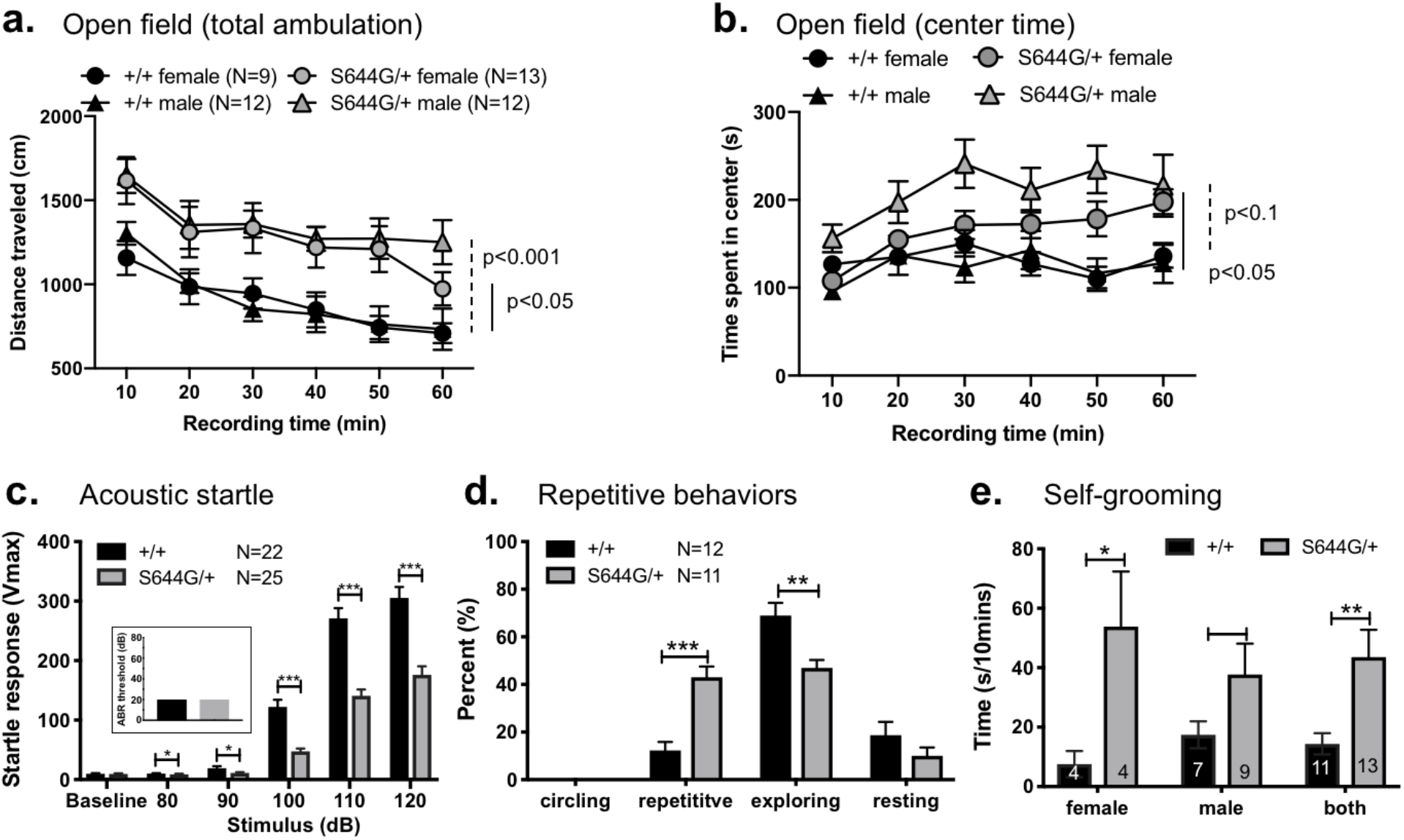
Behavioral phenotypes in B6NJ-*Grin2a*^S644G/+^ mice. (a) Distance traveled in the open field was increased in both sexes of B6NJ-*Grin2a*^S644G/+^ mice, (b) Time spent in the center of the open field was increased in female B6NJ-Grin2a^S644G/+^ mice. *p*-values shown are for the genotype effect in a 2-way repeated measures ANOVA. S644G/+ mice exhibited a significantly decreased acoustic startle response to 90-120 dB stimuli (c) but no hearing impairment as per the ABR threshold test (inset). (d) Increased repetitive behavior (back flipping and grooming events), (e) self-grooming. Error bars indicate S.E.M. *p<0.05; **p<0.01; ***p<0.001; ****p<0.0001, Mann-Whitney rank-sum test.

Heterozygotes of both sexes exhibited a significantly reduced acoustic startle response (**Fig. 4c; Supplementary Fig. 6**) but a normal auditory brainstem response threshold (**Fig. 4c inset**), indicating that hearing *per se* was not impaired but the reaction to stimulus is altered. Because NMDAR mutations are associated with autism spectrum disorder (ASD, (Hacohen *et al*., 2016; Grea *et al*., 2017; Rossi *et al*., 2017) and repetitive behavior in mice (Lee *et al*., 2015; Kim *et al*., 2019), we evaluated additional ASD-relevant behaviors using established criteria in rodents. Although S644G/+ mice showed no impairments in the same-sex reciprocal social interaction test (**Supplementary Fig. 7a-b**), both sexes and strain backgrounds exhibited a significant increase in repetitive behaviors, such as vertical jumping and self-grooming (**Fig. 4d-e; Supplementary Fig. 7c-d**).

Altogether, the impaired maternal care and neurobehavioral features of S644G/+ mice delineate a complex neurological phenotype, with the most prominent features being hyperactivity, decreased anxiety-like behavior, and increased repetitive behaviors.

### NMDARs containing GluN2A-S644G are gain-of-function channels with a slow response time course

We used two-electrode voltage clamp current recordings in *Xenopus laevis* oocytes to evaluate the effect of S644G on NMDAR function. GluN2A-S644G-containing NMDARs significantly increased glutamate potency (**Supplementary Fig. 8a, Table 1**) and glycine potency (**Supplementary Fig. 8b, Table 1**) by 17- and 11-fold, respectively. There was a significant decrease in Zn^2+^ and H^+^ potency, but no detectable change in voltage-dependent Mg^2+^-block (**Supplementary Fig. 8c-e, Table 1**). We evaluated mutation effects on the deactivation time course following rapid removal of glutamate, which controls time course of excitatory postsynaptic currents in the synapse (Lester *et al*., 1990). Prolonged application of the endogenous agonist glutamate (in the presence of glycine) effectively causes channel opening to levels comparable between wildtype and mutant channels. Following a rapid switch from glutamate plus glycine to glycine alone, the time constant (τ) describing deactivation was significantly prolonged in the GluN2A-S644G compared to wildtype (**Fig. 5a-b; Table 1**).

**Figure 5.**
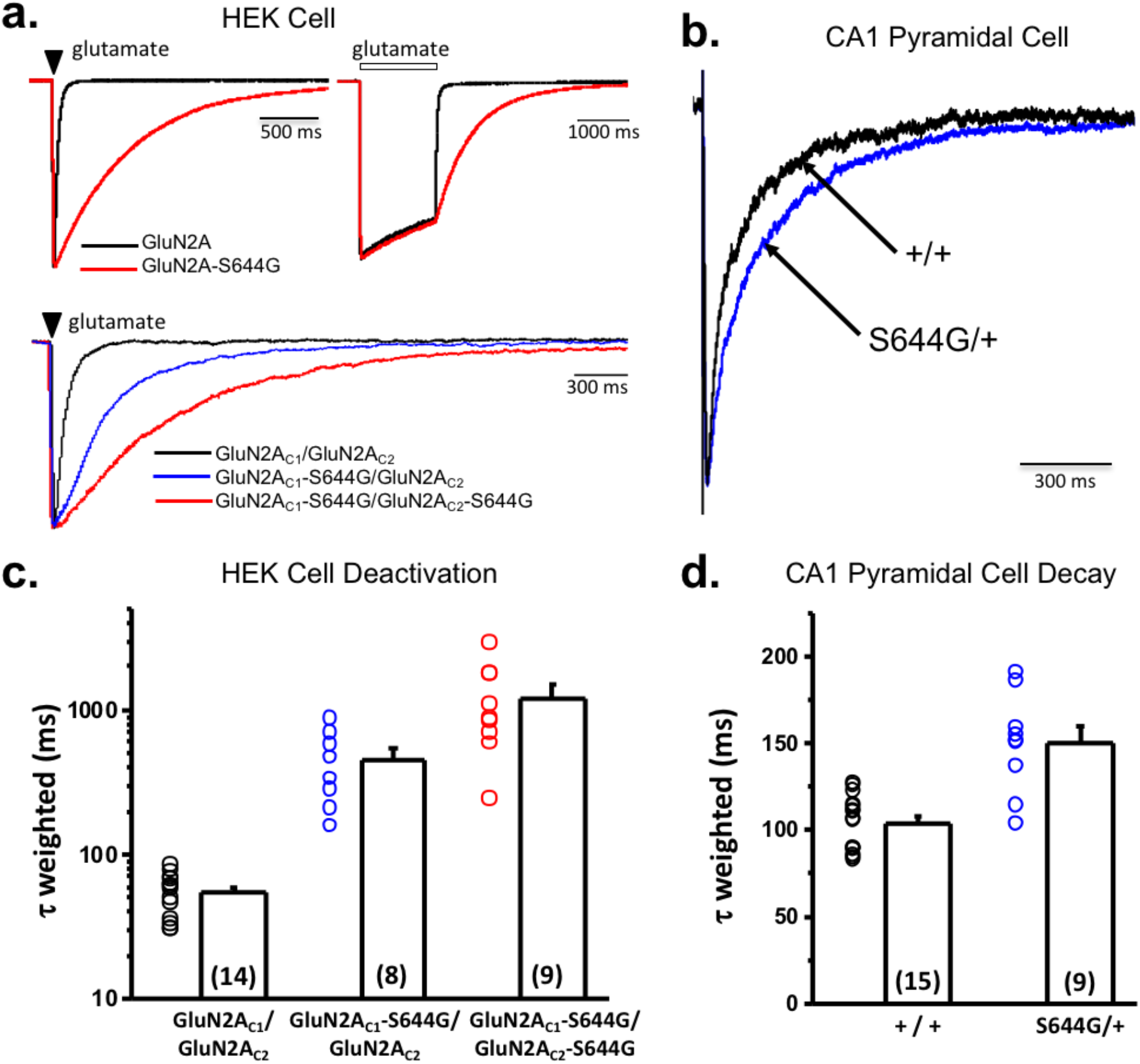
GluN2A-S644G NMDAR response time course *in vitro* and synaptic time course in acute hippocampal slices. (a) *Upper panel* Representative GluNl/GluN2A (black) or GluN1/GluN2A-S644G (red) whole cell current time course in response to brief (1-5 ms, *left panel)* or prolonged (2 s, *right panel)* application of maximally effective concentrations of glutamate (100 μM) and glycine (30 μM). The *lower panel* shows the response for triheteromeric GluN1/GluN2A_c1_/GluN2A_c2_ receptors that contained 0 (black), 1 (gray) or 2 (red) copies of the S644G mutation (brief pulses 20-50 ms; traces corrected for series resistance using ChanneLabv2 software, (Traynelis, 1998); statistics are given in Table 1 and Supplementary Table 3). (b) Superimposed, normalized evoked NMDAR-mediated component of the EPSC onto CA1 pyramidal cells in hippocampal slices from wild +/+ and +/S644G mice. (c) Mean + SEM for the weighted tau describing deactivation for NMDARs expressed in HEK cells. (d) Mean + SEM for the weighted tau describing synaptic decay of the NMDAR-mediated current component of the EPSC onto CA1 pyramidal cells in +/+ or S644G/+ slices. See Supplementary Table 2 and 3 for statistical analysis of all measured parameters. ****p<0.0001, two-sample Student’s |t|-test.

**Table 1:** Functional properties of diheteromeric GluN1/GluN2A-S644G NMDARs

|  | hGluN2A / hGluN2A | hGluN2A-S644G / hGluN2A-S644G |
| --- | --- | --- |
| *Xenopus laevis* oocytes |  |  |
| **Glu, EC<sub>50</sub> (μM)** | 3.0 (2.5, 3.5; 10) | 0.18 (0.15, 0.20; 12) * |
| **Gly, EC<sub>50</sub> (μM)** | 0.86 (0.55, 1.1; 12) | 0.08 (0.047, 0.094; 14) * |
| **Mg<sup>2+</sup>, IC<sub>50</sub> (μM)** | 26 (17, 31; 11) | 33 (27, 35; 26) |
| **pH, IC<sub>50</sub>** | 7.02 (6-17) | 5.86 (8-17) |
| **Current pH 6.8 / pH 7.6 (%)** | 44 ± 1.3 (8) | 99 ± 0.0.7 (8)* |
| **Zn<sup>2+</sup> IC<sub>50</sub> (nM)** | 23 (6.7, 28; 9) | ND |
| **Inhibition by 300nM Zn<sup>2+</sup> (%)** | 56 ± 4.2 (9) | 8.1 ± 1.6 (8)* |
| **P<sub>OPEN</sub> (from MTSEA)** | 0.15 ± 0.01 (13) | 1.0 ± 0.29* (20) |
| *HEK-293 cells* |  |  |
| **Amplitude (peak, pA/pF)** | 70 ± 15 (5) | 194 ± 81 (7) |
| **Amplitude (SS, pA/pF)** | 46 ± 9.1 (5) | 81 ± 27 (7) |
| **Current (SS) / Current (Peak)** | 0.70 ± 0.04 (5) | 0.57 ± 0.13 (7) |
| **10-90% Rise time (ms)** | 9.0 ± 1.2 (5) | 8.5 ± 0.9 (7) |
| **τ<sub>FAST</sub> deactivation (ms)** | 25 ± 6.3 (5) | 1077 ± 292 (7) |
| **τ<sub>SLOW</sub> deactivation (ms)** | 503 ± 223 (5) | 3265 ± 1022 (7) |
| **τ<sub>FAST</sub> amplitude (%)** | 90% (5) | 67% (7) |
| **τ<sub>W</sub> deactivation (ms)** | 48 ± 4.7 (5) | 2129 ± 608** (7) |
| **Charge transfer (pA ms/pF)** | 3,454 (5) | 173,518 (7) |

Potency data are mean (95% CI determined from logEC<sub>50</sub>; n) \* indicates non-overlapping confidence intervals. Agonist concentration-response curves were fitted by:
where EC<sub>50</sub> is the concentration of agonist that produces a half maximal response and *N* is the Hill slope. Proton and Zn<sup>2+</sup> inhibition curves were fitted by
where minimum is the residual response in saturating inhibitor; for proton inhibition, the *minimum* was set to 0.
Other data are mean ± SEM. ND indicates no detectable effect of Zn<sup>2+</sup>. \*\* indicates p<0.05 by unpaired t-test for tau weighted, inhibition by 300 nM Zn<sup>2+</sup>, and inhibition by pH 6.8. Power to detect an effect size of 1.5 for pH, Zn<sup>2+</sup>, and P<sub>open</sub> was >0.8. Power to detect an effect size of 2 for τ<sub>w</sub> (e.g. 20 ms change) was 0.87 (GPower 3.0).

NMDAR open probability depends on subunit composition and varies over 50-fold (Wyllie *et al*., 1998; Erreger *et al*., 2005; Yuan *et al*., 2005; Dravid *et al*., 2008). NMDARs containing GluN1-A652C can be irreversibly locked in the open position upon treatment with the sulfhydryl-modifying reagent methanethiosulfonate ethylammonium (MTSEA), thereby allowing for the evaluation of agonist-induced open probability (Jones *et al*., 2002; Yuan *et al*., 2005; Yuan *et al*., 2009). Co-application of 200 μM MTSEA with glutamate and glycine potentiated the agonist-evoked current for GluN2A-S644G-containing NMDARs compared to wildtype (**Table 1**). Together, these data suggest that the S644G mutation results in a gain-of-function channel with slow kinetics.

Patients with GRIN2A-associated DEE typically have *de novo* heterozygous variants; thus, receptors that contain one wildtype and one variant copy of GluN2A. To investigate whether the NMDARs with one copy of GluN2A-S644G show altered function, we expressed GluN1/GluN2A_c1_/GluN2A_c2_ NMDARs that contained GluN2A subunits modified by addition of coiled-coil domains and an ER retention signal on the C-terminus (Hansen *et al*., 2014) in*Xenopus* oocytes and evaluated the concentration–response curves for glutamate and glycine. GluN1/GluN2A_c1_-S644G/GluN2A_c2_ NMDARs containing one copy of the variant subunit increased the glutamate potency, as measured by reduction in the half-maximally effective concentration of agonist (EC_50_) from 4.7 μM to 0.8 μM. NMDARs that contained two copies of the variant subunit, GluN1/GluN2A_c1_-S644G/GluN2A_c2_-S644G increased the glutamate potency further to 0.09 μM. Glycine potency was shifted in a similar manner as glutamate (**Supplementary Fig. 8H-I; Supplementary Table 1**).

Next, we used whole cell patch clamp recordings to examine the effects of synaptic expression of the GluN2A-S644G subunit in S644G/+ hippocampal brain slices. (Neurons from S644G/S644G homozygous brain slices were not possible to patch). Monosynaptic glutamate release was initiated via Schaffer collateral stimulation, and the NMDAR-mediated current component was isolated at −30 mV in 0.2 Mg^2+^ via bath application of AMPAR and GABAAR antagonists (see Methods). Although there was no difference in the peak NMDAR current response (**Supplementary Table 2**), weighted tau, a measurement of receptor deactivation, was significantly increased in S644G/+ mice (**Fig. 5b, d; Table 1**). There were no detectable changes in other measurements of NMDAR time course between +/+ and S644G/+ mice (**Supplementary Table 2**), and no detectable difference in membrane resistance, series resistance, or cell capacitance, or APV-sensitive leak current (data not shown, see Methods). Similarly, utilizing the triheteromeric NMDAR expression system as described above, we determined the effect of 0, 1 and 2 copies of GluN2A-S644G subunits on weighted tau (**τ_W_**) in NMDARs expressed in HEK-293 cells. Our results suggest that **τ_W_** was significantly different for 1 or 2 copies of GluN2A-S644G compared to wildtype GluN2A (p<0.05, one-way ANOVA, Tukey’s multiple comparison test, **Supplementary Table 3** and Fig. 5a, lower panel), suggesting that our whole cell patch data from hippocampal slice experiments might have shown modest difference between wild type and heterozygous animals due to the presence of different triheteromeric NMDAR combinations. These results indicate that the synaptic NMDAR-mediated current component is significantly enhanced in S644G/+ mice, in agreement with heterologous expression data.

### Hyperexcitability and abnormal network activity of *Grin2a^S644G^* primary neurons

We examined effect of S644G on neuronal network hyperexcitability using multielectrode array (MEA) recording of primary cortical neurons derived from neonatal mice, recorded every other day from five days *in vitro* (DIV5) to DIV29.

Early timepoints displayed expected low spontaneous activity and minimal network events (spikes and bursts) for all genotypes. By DIV13, networks from all genotypes had reached the maximum number of active electrodes (nAE) per well, indicating robust development of network activity (**Supplementary Fig. 9a**). Both S644G/+ and S644G/S644G networks displayed significantly increased mean spontaneous firing rate compared to wildtype (**Fig. 6a**). Mutant networks also had an increased burst rate relative to wildtype (**Fig. 6b**)

**Figure 6.**
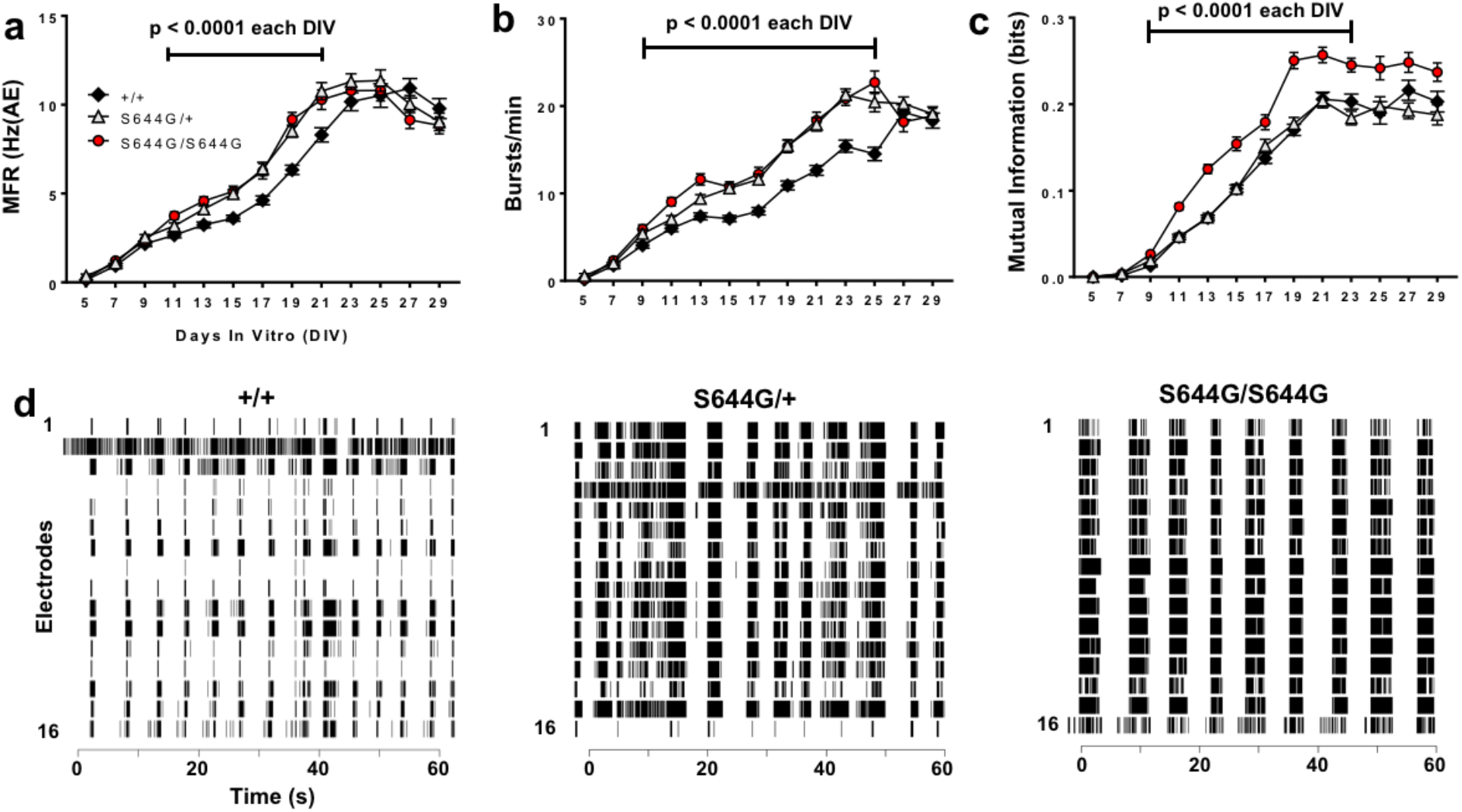
Multielectrode array (MEA) network phenotypes. We analyzed the activity of cortical neural networks from six litters of wildtype (+/+, n=108 wells) and S644G/+ (n=107 wells), and 5 litters of S644G/S644G (n=80 wells) mice. (a) Mutant neurons exhibit significantly elevated mean firing rate (MFR; expressed relative to the number of active electrodes (AE)). (b) Both mutant genotypes have a significantly higher burst rate compared to wildtype. (c) S644G/S644G networks display an increased local synchrony compared to both S644G/+ and wildtype. Error bars in a-c indicate SEM. (d) Representative Raster plots of network activity at DIV21 for each genotype. For statistical analysis, features from DIV9-DIV29 were each rank- and then normal quantile-transformed and fit in a least-squares regression model using genotype and plate as covariates. The *p*-values obtained for genotype effect were adjusted using a Bonferroni correction for the 11 DIVs analyzed for each feature. For genotype effects significant (*p*<0.05) after Bonferroni correction, a post-hoc Dunne’s test was used (alpha=0.05) to examine the effect of heterozygosity or homozygosity with wildtype as control. Supplementary Table 5 shows the genotype effect and plate effect *p*-values.

Based on the altered bursting properties of mutant networks, we examined whether they also exhibit aberrant coordinated activity. All genotypes displayed the emergence of network bursts, which are synchronized periods of activity followed by quiescent periods (Chiappalone *et al*., 2006; Wagenaar *et al*., 2006) (Van Pelt *et al*., 2004; Vajda *et al*., 2008; Cotterill *et al*., 2016); however, both heterozygous and homozygous networks had a significantly longer network bursts relative to wildtype (**Supplementary Fig. 9c**).

We investigated synchronous network firing using the mutual information algorithm, a pairwise cross-correlation of electrode activity among neighboring electrodes that gives an indication of spatial connectivity (Gelfman *et al*., 2018). Mutual information analyses revealed elevated synchrony of S644G/S644G networks relative to S644G/+ or wildtype (**Fig 6c**). Collectively, these data suggest that S644G/S644G networks have altered network topology compared to S644G/+ or wildtype. Representative raster plots highlight and summarize the network features of each genotype (**Fig. 6d**).

To assess activity stimulation via MEA, we treated wildtype and mutant networks with NMDA as it mimics glutamate action while being solely selective for NMDA receptors. As expected mutant networks treated with NMDA demonstrated clearly enhanced spontaneous activity differences dependent on gene dosage such that S644G/S644G networks exhibited significantly higher firing rates than S644G/+ networks, which were themselves significantly more sensitive than wildtype (**Supplementary Fig. 9d**). Activity declined in all genotypes at higher doses of NMDA with mutants being more strongly affected. Network responses to agonist were further probed using electrical stimulation (see **Methods**). Electrical stimulation in the presence of exogenous NMDA resulted in sustained activity of S644G/S644G networks compared to wildtype (**Supplementary Fig. 9e**). Administration of NMDA to wildtype networks did not result in an increased evoked response duration (**Supplementary Fig. 9e**) further demonstrating that homozygous networks possess enhanced susceptibility to agonist, which likely causes the mutant hyperactivity and altered bursting properties. Overall, our MEA studies suggest that the seizure phenotypes observed *in vivo* correlate with robust, quantifiable phenotypes in cultured neuronal networks.

### Prospects of NMDAR antagonist combination therapy

The initial seizure frequency in the *GRIN2A* S644G patient was approximately 200 per month, with no seizure-free days (**Supplementary Fig. 10a**). Prior to a genetic diagnosis, empirical treatment trials with vigabatrin, topiramate, levetiracetam, and clobazam were unsuccessful in reducing seizures. After the genetic diagnosis, treatment with the NMDAR antagonist memantine was considered as a rational candidate because of reports of this agent’s efficacy and safety in children with other *GRIN2A* gain-of-function variants (Pierson *et al*., 2014) (Hosenbocus and Chahal, 2013; Nguyen *et al*., 2016). Given our functional data supporting a gain-of-function effect and the effectiveness of memantine *in vitro* (see below and **Supplementary Table 3-4**), treatment with memantine began at age 2 years and reduced the daily seizure burden by half. This was followed by treatment with dextromethorphan (DM), also used safely for other indications in children, at age 3 years at doses of 5 mg/kg/day increasing to 10 mg/kg/day. While there was not substantial additional benefit, we observed some worsening of seizures with attempts to wean DM. The patient, now 6 years old, experiences 3-5 brief tonic seizures and 1-2 myoclonic seizures per day on a combination of memantine, DM, and zonisamide (**Supplementary Fig. 10**). Profound intellectual disability, spastic quadriparesis, cortical visual impairment and oropharyngeal dysphasia persist.

Exploiting the viability of *Xenopus laevis* oocytes expressing the mutant channel, NMDAR drug candidates were tested and efficacy in half-maximal inhibitory responses were assessed. Different classes of NMDAR antagonists inhibited channel activation with modestly reduced potency compared to wild type receptors (**Supplementary Fig. 8 f,g**). Because neurons express receptors that contain one copy of the mutant allele, as well as tri-heteromeric receptors that contain mutant GluN2A and GluN2B, we expressed receptors in which we could control subunit stoichiometry using coiled-coil domain masking of an ER-retention signal. **Supplementary Fig. 8** shows the properties of the receptors, and **Fig. 7a-b** shows the potency of two FDA-approved drugs (memantine and DM). The inhibitory effects of memantine and DM were corroborated in cortical networks on MEA (**Fig. 7c-d**). Together, these data encouraged us to test whether NMDAR antagonists would mitigate lethal seizures of S644G homozygous pups.

**Figure 7.**
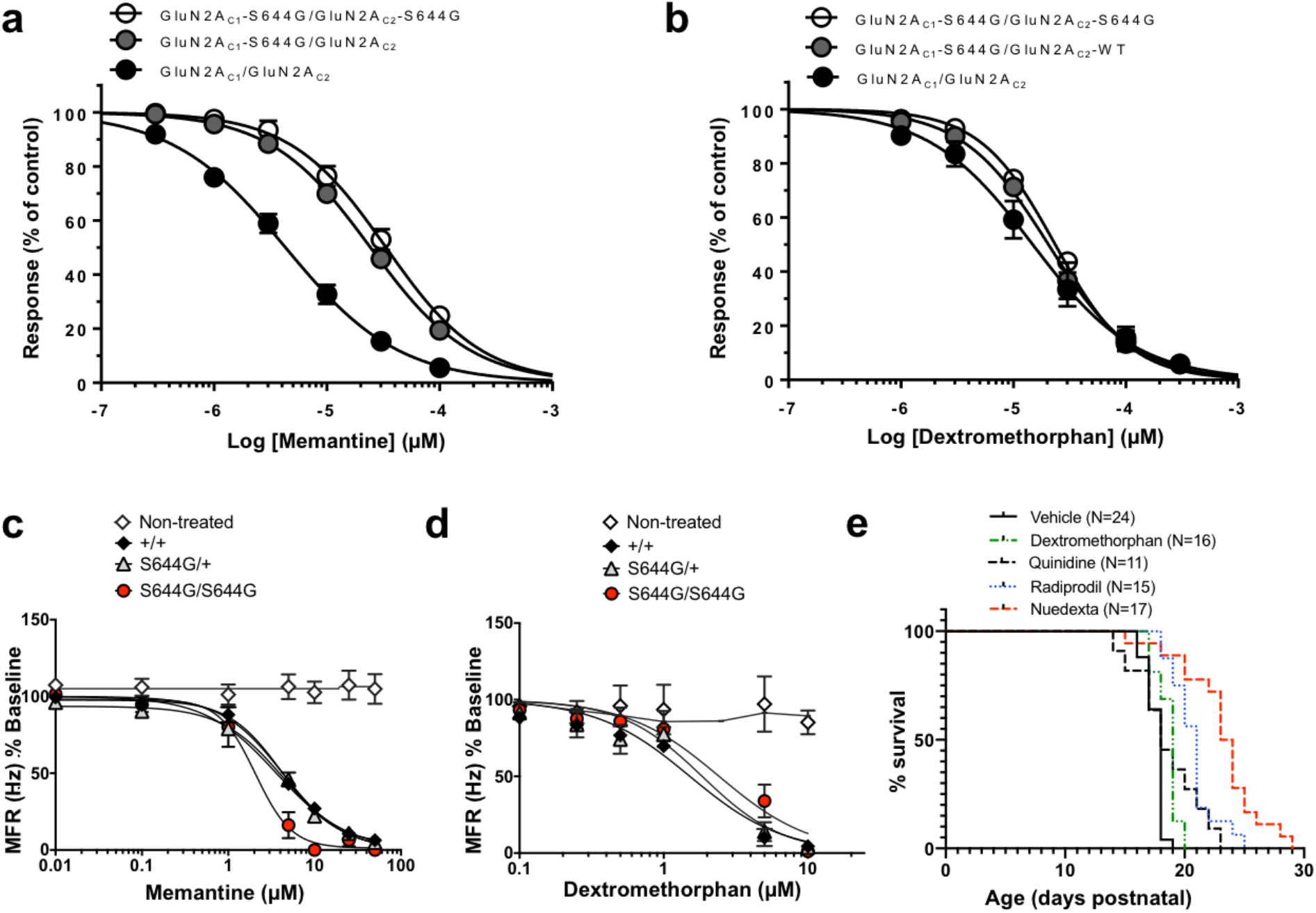
*In vitro* and *in vivo* drug responses – impact on network hyperexcitability and lethal seizures. (a, b) Composite concentration-response curves for memantine (n=1-12 oocytes) and dextromethorphan (n=10-13) on NMDA receptors that contained 0, 1, or 2 mutant GluN2A-S644G subunits. The IC_50_ values for single and double copy GluN2A-S644G mutant receptors for both dextromethorphan and memantine were significantly different from WT-GluN2A receptors (p<0.05, ANOVA and Neuman-Kuels multiple comparison test; Supplementary Table 4). (c,d) Fitted concentration-response curves for memantine and dextromethorphan on MEA. Data for each well are normalized to baseline firing collected before treatment. Memantine; n=3 litters for wildtype and S644G+ (≥ 21 wells each concentration), n=1 litter for S644G/S644G (4 wells each concentration). Dextromethorphan; n=5 litters for wildtype and S644G/+ (≥ 22 wells each concentration), and 2 litters for S644G/S644G (12 wells each concentration). Non-treated controls; 42 and 24 wells memantine and dextromethorphan, respectively. Compound IC_50_ values are shown on Supplementary Table 6. Genotype x treatment interaction effects were tested in a least-squares regression model of rank- and then normal quantile-transformed data. (e) Pharmacological rescue of lethal seizures of S644G/S644G homozygotes, showing the respective survival after daily injections of dextromethorphan (DM), quinidine, radiprodil, and Nuedexta, as discussed in the text. The respective doses were chosen to be just under known toxic doses in mice. The Mantel-Cox log-rank test was used to determine significant differences between curves: vehicle vs. DM, (+ 1 day median survival, χ2=13.8, p<0.0002); vehicle vs. radiprodil, (+ 3 days, χ2=29.1, pO.OOOl); vehicle vs. Nuedexta, (+ 5.5 days, χ2=29.8, p0.0001); vehicle vs. quinidine (+ 1 day median survival, χ2=3.69, p=0.055).

In a first trial, homozygous pups were treated with DM from PND8 to PND21 at the established ED50 of 25 mg/kg (Kim *et al*., 2003). DM extended the survival of homozygous pups by a 1-day median (**Fig. 7e**; p=0.002, Mantel-Cox log-rank test). To improve the exposure of DM, we tried Nuedexta, a combination drug comprised of DM (25 mg/kg) and quinidine (10 mg/kg), which at low concentrations is known to block the rapid metabolism of DM to dextrorphan, a more potent but short-lived metabolite (Pioro *et al*., 2010). Nuedexta treatment of *Grin2a*^S644G^ homozygotes extended pup survival significantly further than DM alone, by a median of 5.5 days (**Fig. 7e**; p<0.0001, Mantel-Cox log-rank test). Conversely, quinidine alone only modestly extended survival by a median of 1 day (**Fig. 7e** p<0.02, Mantel-Cox log-rank test). Because *Grin2a*^S644G^ show overactive NMDAR currents, we also tested whether radiprodil (2 mg/kg) prolongs survival, as it blocks a different population of GluN2B-containing NMDARs (Borza and Domany, 2006; Mony *et al*., 2009). Radiprodil extended survival by 3 days – further than DM alone, but not as long as Nuedexta, (**Fig. 7e**; p<0.0001, Mantel-Cox log-rank test). Although the severe seizures of S644G homozygotes do offer a very robust challenge for testing therapeutic intervention, these results support further investigation into NMDAR antagonist therapies.

## DISCUSSION

The *GRIN2A* S644G substitution in a patient with DEE, on which our studies were based, is situated in the highly conserved signature gating sequence (SYTANLAAF) in TM3, the third transmembrane pore region of the NMDAR. Amongst all variants we have examined, it displays one of the strongest gain-of-function effects on NMDAR activity. *Grin2a*^S644G^ mutant mice were developed as part of a larger effort to model the pathophysiologic features of children with DEE due to gain-of-function *GRIN2A* variants, and in which to further explore therapies. The key features of GRIN2A-associated DEE are medically refractory epilepsy and developmental delay/intellectual disability. *Grin2a*^S644G^ mice mimic several aspects of human disease, including seizure susceptibility and distinct and robust, if not unusual behavioral phenotypes.

Interestingly, *Grin2a*^S644G^ mice are both seizure susceptible (spontaneous lethal seizures) and resistant (induced partial seizures), with each being a robust, fully penetrant trait. Given the breadth and extent of *Grin2a* expression in the brain, we strongly suspect these differences arise from pleiotropy in different neuron types and brain regions, as has been suggested or shown directly with the advent of conditional mutations in other mouse seizure models (Papale *et al*., 2009; Wagnon *et al*., 2011; Asinof *et al*., 2015; Makinson *et al*., 2016). Pleiotropy is not unexpected for widely and highly expressed genes such as *Grin2a* and likely also contributes to the diversity of comorbid behaviors, as shown previously for *Dnm1*^Ftfl^, the mouse model of *DNM1* EE (Asinof *et al*., 2015).

NMDAR gene variants, including those in *GRIN2A*, are associated with autism spectrum disorder (Yuan *et al*., 2015) and aphasia (Carvill *et al*., 2013), and *GRIN2A* has recently been implicated in ADHD (Slopien *et al*., 2006). The features in *Grin2a*^S644G^ heterozygous mice of hyperactivity, decreased anxiety, and a variety of stereotypies are consistent with the human disease spectrum (Hu *et al*., 2016). Notably, in our patient and in many with DEE, the level of cognitive function is sufficiently low to preclude reliable anxiety measurements. We also note that our patient with *GRIN2A* S644G showed no evidence of hyperactivity or stereotypies. However, in many genetic DEEs there can be significant phenotypic heterogeneity among patients with the same primary gene defect – even when harboring the identical variant (Helbig and Tayoun, 2016) – which, when also considering the many genetic differences across and within species, warrants caution about attempting to cross-correlate phenotypes too narrowly.

Nevertheless, rodent models do offer unique advantages to probe vulnerabilities at the circuit and cellular level caused by the orthologous genetic defect. Specifically, the relative resistance of *Grin2a*^S644G^ heterozygotes to 6 Hz partial seizures suggests altered limbic system function or connectivity (Barton *et al*., 2001; Metcalf *et al*., 2017). Ironically, vulnerability in the limbic system is also suggested by the lethal seizures whose features include a wild run followed rapidly by tonic-clonic episodes and tonic hind limb extension, resembling audiogenic seizures which are known to involve paroxysmal activity in the cortex, medial geniculate, hippocampus, and amygdala (Marescaux *et al*., 1987; Naritoku *et al*., 1992; Dutra Moraes *et al*., 2000; Faingold, 2012); the main structures of limbic circuitry (Giordano *et al*., 2015; Giordano *et al*., 2016)

Functional characterization of mutant channel activity shows a very strong gain-of-function effect and slower kinetics, supported by assessment of synaptic activity in excitatory neurons of mutant mice. Evaluation of receptors with 0, 1, or 2 copies of the mutant allele revealed an intermediate effect of S644G when only one copy of the variant is present in the complex. Although the synaptic data indicate a prolongation of NMDAR-mediated current in the mutants, the results – while significant – were less striking than in heterologous assays, even for the intermediate phenotype. Several potential explanations for this exist, including likely binomial probability of receptors having receptors with 0, 1, or 2 copies of the mutant, and wildtype subunits will confer faster synaptic time course. In addition, some receptors at this age will contain two copies of GluN2B in the synapse. Tri-heteromeric NMDARs that contain one GluN2B subunit and one GluN2A-S644G subunit may also reside in the synapse since both GluN2A and GluN2B expression are high in excitatory pyramidal cells in developing mice (Monyer *et al*., 1994; Hansen *et al*., 2014). There also may be systematic bias towards healthy neurons that allowed stable patch clamp recording, which may have selected against cells with strong expression of GluN2A-S644G.

MEA examination of activity patterns in mutant primary neurons and their longitudinal study during network establishment revealed the development of hyperactive mutant networks. S644G heterozygotes and homozygotes have similar mean firing rates and bursting features, perhaps reflecting more cell-autonomous effects, but homozygotes stand-out in more complex measures of network synchronicity.

The presence of quantifiable excitability features and the mirroring of pharmacological rescue of aberrant activity by NMDAR antagonists is also validated on the MEA platform, representing a significant opportunity for preclinical intervention screening prior to laborious *in vivo* testing. Screening therapeutic compounds in both *in vitro* and *ex vivo* platforms should identify compounds with better efficacy than in either alone. The prominent lethal seizures of homozygous mutants also afford a robust challenge to explore the potential of NMDAR antagonists as therapeutic agents. Although the homozygous S644G mice eventually succumbed to lethal seizures irrespective of treatment regimen, the extension of survival by 5.5 days (or 30% of their lifespan) using Nuedexta is a significant indication that further studies of NMDAR targeted therapies are warranted.

These combined results of a DEE variant in a patient, heterologous expression systems, a mouse model, and murine neurons in culture empower a range of experiments to capture the core phenotypes of epilepsy and developmental disability. Similar phenotypes of DEE are observed in patients with other *GRIN2A* variants, and gain of function has been demonstrated using the oocyte system used in this report. Together with application of targeted pharmaceutical and genetic approaches to therapy, we anticipate that our observations for *GRIN2A* S644G will have relevance to the broad group of patients with gain-of-function NMDAR variants.

## ACKNOWLEDGMENTS

We are grateful to our patient’s family for participating in our research to allow for sharing of deidentified data. We would like to thank Sophie Colombo and Sahar Gelfman for helpful discussions regarding MEA analyses, and Aamir Zuberi for his help in generating the mouse model.

## SUPPLEMENTARY FIGURE LEGENDS

Supplementary Figure 1. Genetic transmission of S644G allele and husbandry of *Grin2a*
^S644G^ mice. Mendelian distribution of +/+, S644G/+, and S644G/S644G genotypes in (a) B6NJ and (b) F1 or F2 hybrid background. Observation gathered from multiple litters (>50). (c) Pups born from B6NJ S644G/+ females show no milk spot, indicative of poor maternal care.

Supplementary Figure 2. Representative wake EEG of *Grin2a* S644G/+ and wildtype +/+ littermates. No obvious epileptiform activity was observed in traces from at least seven *Grin2a*^S644G/+^ mice recorded for 48 hours each.

Supplementary Figure 3. GluN2A, GluN2B protein and mRNA in whole brain of two week old mouse pups. (a) Representative western blot of whole brain lysates probed for GluN2B, GluN2A, PSD95, β3-tubulin. (b) Plot of amount of protein normalized to β3-tubulin and wildtype. (c) mRNA expression quantification of total GRIN2B, GRIN2A, and PSD95 was determined by qRT-PCR and normalized to housekeeping gene, GADPH. Error bars indicate S.E.M. *p<0.05, **, ***, **** Student’s |t|-test

Supplementary Figure 4. Additional open field parameters – Ambulation (a) and center time (b) and vertical activity in B6NJ and F1 hybrid mice comparing S644G/+ and +/+ (wildtype) genotypes. *p*-values shown are for the genotype effect in a 2-way repeated measures ANOVA.

Supplementary Figure 5. Elevated plus maze. Elevated plus maze test was performed of S644G/+ and +/+ mice on both B6NJ and F1 hybrid strain backgrounds, both sexes combined, showing the # of open arm entries (a), the percent time spent on the open arms (b), vs. total entries and time spent at the junction of the arms. There was a trend for S644G/+ mice to spend more time on the open arms (p=0.051, Mann-Whitney rank-sum test, strain backgrounds combined).

Supplementary Figure 6. a. Acoustic startle response (F1 hybrid), b. representative ABR traces

As in the B6NJ line, heterozygous mice on the F1 background exhibited a markedly decreased response to a range of acoustic startle stimuli, without displaying impairments in the ABR test, indicating that reactivity to acoustic stimuli was altered, independent of defects in sensory perception.

Supplementary Figure 7. a, b. Social interaction (B6NJ and F1 hybrid), c, d. repetitive behaviors (F1 hybrid)

As in the B6NJ line, heterozygous mice on the F1 background exhibited normal social interaction behaviors and significantly increase repetitive behaviors, especially self-grooming.

Supplementary Figure 8. Response of S644G-containing NMDARs to agonists *in vitro*. (a, b) Fitted composite glutamate and glycine concentration-response curves for diheteromeric human wildtype GluN1/GluN2A- and GluN1/GluN2A-S644G expressed in *Xenopus laevis* oocytes. Two-electrode voltage-clamp recordings were conducted at a holding potential of −40 mV. Glutamate potency was quantified as the half-maximally effective response (EC_50_) of recombinant NMDARs receptors comprised of wild type or mutant GluNR2A in the presence of a maximally effective concentration of glycine (100 μM).Glycine potency was evaluated in the presence of a maximally effective concentration of glutamate (100 μM). (c, d) Fitted concentration-response curves for endogenous antagonists, Mg^2+^ (at −60 mV) and Zn^2+^(at −20 mV) at diheteromeric human GluN1/GluN2A NMDARs. (e,f) Fitted concentration-response curves for FDA-approved drugs (dextromethorphan, memantine) at diheteromeric human NMDARs. (g) Percentage current response at pH6.8 versus 7.6 shows decreased current attenuation in the presence of increased proton concentrations (ie, low pH). Concentration-response curves were generated for co-agonists glutamate and glycine, channel blocker Mg^2+^ and endogenous antagonist Zn^2+^ to evaluate whether the mutation changes agonist potency or the sensitivity to endogenous modulators. (h) Agonist potency shifts by GluN2A-S644G-containing NMDARs *in vitro*. Fitted concentration-response curves for glutamate, in the presence of 100 μM glycine (a), and for glycine in the presence of 100 μM glutamate (i) for NMDARs that contain 0, 1, or 2 copies of the rat GluN2A-S644G variant expressed in *Xenopus laevis* oocytes. Results are mean + SEM from 12-17 (glutamate) or 13-14 oocytes (glycine) from two separate injections. Either 1 or 2 copies of theGluN2A-S644G mutation significantly increases both glutamate and glycine potency of the NMDA receptors (p<0.05, ANOVA, Neuman-Kuels multiple comparison test).

Supplementary Figure 9. Additional multielectrode array analyses

(a) Temporal development of number of active electrodes, defined as electrodes recording at least 5 spikes per minute, demonstrates >94% active electrodes for each genotype by DIV13. (b) Mutant networks displayed significantly longer network bursts (NB) relative to wildtype networks. Error bars in (a) and (b) indicate SEM – statistical tests were done as described in Figure 6 and results listed on Supplementary Table 4. (c) Concentration-response curve of NMDA demonstrates sensitivity of mutant networks to NMDA-mediated excitation. Data for each well are normalized to the baseline firing recorded prior to agonist addition. Data were derived from 4 independent cultures with ≥17 wells for each genotype. (d) Peristimulus time histograms (400 ms) (see Materials and Methods) pre- and post-exposure to specified NMDA concentrations (n=6 wells). At baseline, mutant networks display prolonged evoked burst duration compared to wildtype with a mean AUC of 450 ± 10.7 compared to 330 ± 4.9 for wildtype. NMDA addition decreased evoked burst duration for both wildtype and mutant neurons in a concentration-dependent fashion. 1 μM NMDA; wildtype AUC 300 ± 5.4, S644G/S644G AUC 340 ± 8.8. 10 μM NMDA; wildtype AUC 250 ± 6.2 and S644G/S644G 140 ± 4.7.

Supplementary Figure 10. Pharmacotherapy in the patient with GRIN2A S644G. Seizure frequency is shown during experimental treatment paradigm during which time the antiepileptic medications for the patient were altered.

## REFERENCES

Akazawa C, Shigemoto R, Bessho Y, Nakanishi S, Mizuno N. Differential expression of five N-methyl-D-aspartate receptor subunit mRNAs in the cerebellum of developing and adult rats. J Comp Neurol 1994; 347(1): 150–60.

Asinof S, Mahaffey C, Beyer B, Frankel WN, Boumil R. Dynamin 1 isoform roles in a mouse model of severe childhood epileptic encephalopathy. Neurobiol Dis 2016; 95: 1–11.

Asinof SK, Sukoff Rizzo SJ, Buckley AR, Beyer BJ, Letts VA, Frankel WN, et al. Independent Neuronal Origin of Seizures and Behavioral Comorbidities in an Animal Model of a Severe Childhood Genetic Epileptic Encephalopathy. PLoS Genet 2015; 11(6): e1005347.

Bar-Shira O, Maor R, Chechik G. Gene Expression Switching of Receptor Subunits in Human Brain Development. PLoS Comput Biol 2015; 11(12): e1004559.

Barton ME, Klein BD, Wolf HH, White HS. Pharmacological characterization of the 6 Hz psychomotor seizure model of partial epilepsy. Epilepsy Res 2001; 47(3): 217–27.

Borza I, Domany G. NR2B selective NMDA antagonists: the evolution of the ifenprodil-type pharmacophore. Curr Top Med Chem 2006; 6(7): 687–95.

Carvill GL, Regan BM, Yendle SC, O’Roak BJ, Lozovaya N, Bruneau N, et al. GRIN2A mutations cause epilepsy-aphasia spectrum disorders. Nat Genet 2013; 45(9): 1073–6.

Chen C, Okayama H. High-efficiency transformation of mammalian cells by plasmid DNA. Mol Cell Biol 1987; 7(8): 2745–52.

Chen W, Tankovic A, Burger PB, Kusumoto H, Traynelis SF, Yuan H. Functional Evaluation of a De Novo GRIN2A Mutation Identified in a Patient with Profound Global Developmental Delay and Refractory Epilepsy. Mol Pharmacol 2017; 91(4): 317–30.

Chiappalone M, Bove M, Vato A, Tedesco M, Martinoia S. Dissociated cortical networks show spontaneously correlated activity patterns during in vitro development. Brain Res 2006; 1093(1): 41–53.

Cotterill E, Charlesworth P, Thomas CW, Paulsen O, Eglen SJ. A comparison of computational methods for detecting bursts in neuronal spike trains and their application to human stem cell-derived neuronal networks. J Neurophysiol 2016; 116(2): 306–21.

de Ligt J, Willemsen MH, van Bon BW, Kleefstra T, Yntema HG, Kroes T, et al. Diagnostic exome sequencing in persons with severe intellectual disability. N Engl J Med 2012; 367(20): 1921–9.

Dravid SM, Prakash A, Traynelis SF. Activation of recombinant NR1/NR2C NMDA receptors. J Physiol 2008; 586(18): 4425–39.

Dutra Moraes MF, Galvis-Alonso OY, Garcia-Cairasco N. Audiogenic kindling in the Wistar rat: a potential model for recruitment of limbic structures. Epilepsy Res 2000; 39(3): 251–9.

Endele S, Rosenberger G, Geider K, Popp B, Tamer C, Stefanova I, et al. Mutations in GRIN2A and GRIN2B encoding regulatory subunits of NMDA receptors cause variable neurodevelopmental phenotypes. Nat Genet 2010; 42(11): 1021–6.

Erreger K, Dravid SM, Banke TG, Wyllie DJ, Traynelis SF. Subunit-specific gating controls rat NR1/NR2A and NR1/NR2B NMDA channel kinetics and synaptic signalling profiles. J Physiol 2005; 563(Pt 2): 345–58.

Faingold CL. Brainstem Networks: Reticulo-Cortical Synchronization in Generalized Convulsive Seizures. In: th, Noebels JL, Avoli M, Rogawski MA, Olsen RW, Delgado-Escueta AV, editors. Jasper’s Basic Mechanisms of the Epilepsies. Bethesda (MD); 2012.

Gelfman S, Wang Q, Lu YF, Hall D, Bostick CD, Dhindsa R, et al. meaRtools: An R package for the analysis of neuronal networks recorded on microelectrode arrays. PLoS Comput Biol 2018; 14(10): e1006506.

Giordano C, Costa AM, Lucchi C, Leo G, Brunel L, Fehrentz JA, et al. Progressive Seizure Aggravation in the Repeated 6-Hz Corneal Stimulation Model Is Accompanied by Marked Increase in Hippocampal p-ERK1/2 Immunoreactivity in Neurons. Front Cell Neurosci 2016; 10: 281.

Giordano C, Vinet J, Curia G, Biagini G. Repeated 6-Hz Corneal Stimulation Progressively Increases FosB/DeltaFosB Levels in the Lateral Amygdala and Induces Seizure Generalization to the Hippocampus. PLoS One 2015; 10(11): e0141221.

Grea H, Scheid I, Gaman A, Rogemond V, Gillet S, Honnorat J, et al. Clinical and autoimmune features of a patient with autism spectrum disorder seropositive for anti-NMDA-receptor autoantibody. Dialogues Clin Neurosci 2017; 19(1): 65–70.

Hacohen Y, Wright S, Gadian J, Vincent A, Lim M, Wassmer E, et al. N-methyl-d-aspartate (NMDA) receptor antibodies encephalitis mimicking an autistic regression. Dev Med Child Neurol 2016; 58(10): 1092–4.

Hamdan FF, Gauthier J, Araki Y, Lin DT, Yoshizawa Y, Higashi K, et al. Excess of de novo deleterious mutations in genes associated with glutamatergic systems in nonsyndromic intellectual disability. Am J Hum Genet 2011; 88(3): 306–16.

Hansen KB, Ogden KK, Yuan H, Traynelis SF. Distinct functional and pharmacological properties of Triheteromeric GluN1/GluN2A/GluN2B NMDA receptors. Neuron 2014; 81(5): 1084–96.

Helbig I, Tayoun AA. Understanding Genotypes and Phenotypes in Epileptic Encephalopathies. Mol Syndromol 2016; 7(4): 172–81.

Hosenbocus S, Chahal R. Memantine: a review of possible uses in child and adolescent psychiatry. J Can Acad Child Adolesc Psychiatry 2013; 22(2): 166–71.

Hu C, Chen W, Myers SJ, Yuan H, Traynelis SF. Human GRIN2B variants in neurodevelopmental disorders. J Pharmacol Sci 2016; 132(2): 115–21.

Johnstone AF, Gross GW, Weiss DG, Schroeder OH, Gramowski A, Shafer TJ. Microelectrode arrays: a physiologically based neurotoxicity testing platform for the 21st century. Neurotoxicology 2010; 31(4): 331–50.

Jones KS, VanDongen HM, VanDongen AM. The NMDA receptor M3 segment is a conserved transduction element coupling ligand binding to channel opening. J Neurosci 2002; 22(6): 2044–53.

Kim HC, Shin CY, Seo DO, Jhoo JH, Jhoo WK, Kim WK, et al. New morphinan derivatives with negligible psychotropic effects attenuate convulsions induced by maximal electroshock in mice. Life Sci 2003; 72(16): 1883–95.

Kim S, Kim DG, Gonzales EL, Mabunga DFN, Shin D, Jeon SJ, et al. Effects of Intraperitoneal N-methyl-D-aspartate (NMDA) Administration on Nociceptive/Repetitive Behaviors in Juvenile Mice. Biomol Ther (Seoul) 2019; 27(2): 168–77.

Lee EJ, Lee H, Huang TN, Chung C, Shin W, Kim K, et al. Trans-synaptic zinc mobilization improves social interaction in two mouse models of autism through NMDAR activation. Nat Commun 2015; 6: 7168.

Lesca G, Rudolf G, Bruneau N, Lozovaya N, Labalme A, Boutry-Kryza N, et al. GRIN2A mutations in acquired epileptic aphasia and related childhood focal epilepsies and encephalopathies with speech and language dysfunction. Nat Genet 2013; 45(9): 1061–6.

Lester RA, Clements JD, Westbrook GL, Jahr CE. Channel kinetics determine the time course of NMDA receptor-mediated synaptic currents. Nature 1990; 346(6284): 565–7.

Makinson CD, Dutt K, Lin F, Papale LA, Shankar A, Barela AJ, et al. An Scn1a epilepsy mutation in Scn8a alters seizure susceptibility and behavior. Exp Neurol 2016; 275 Pt 1: 46–58.

Marescaux C, Vergnes M, Kiesmann M, Depaulis A, Micheletti G, Warter JM. Kindling of audiogenic seizures in Wistar rats: an EEG study. Exp Neurol 1987; 97(1): 160–8.

Metcalf CS, West PJ, Thomson KE, Edwards SF, Smith MD, White HS, et al. Development and pharmacologic characterization of the rat 6 Hz model of partial seizures. Epilepsia 2017; 58(6): 1073–84.

Mony L, Kew JN, Gunthorpe MJ, Paoletti P. Allosteric modulators of NR2B-containing NMDA receptors: molecular mechanisms and therapeutic potential. Br J Pharmacol 2009; 157(8): 1301–17.

Monyer H, Burnashev N, Laurie DJ, Sakmann B, Seeburg PH. Developmental and regional expression in the rat brain and functional properties of four NMDA receptors. Neuron 1994; 12(3): 529–40.

Naritoku DK, Mecozzi LB, Aiello MT, Faingold CL. Repetition of audiogenic seizures in genetically epilepsy-prone rats induces cortical epileptiform activity and additional seizure behaviors. Exp Neurol 1992; 115(3): 317–24.

Nguyen L, Thomas KL, Lucke-Wold BP, Cavendish JZ, Crowe MS, Matsumoto RR. Dextromethorphan: An update on its utility for neurological and neuropsychiatric disorders. Pharmacol Ther 2016; 159: 1–22.

Ogden KK, Chen W, Swanger SA, McDaniel MJ, Fan LZ, Hu C, et al. Molecular Mechanism of Disease-Associated Mutations in the Pre-M1 Helix of NMDA Receptors and Potential Rescue Pharmacology. PLoS Genet 2017; 13(1): e1006536.

Otso N. A threshold selection method from gray-level histograms. Automatica 1975; 11: 23–7.

Paoletti P, Bellone C, Zhou Q. NMDA receptor subunit diversity: impact on receptor properties, synaptic plasticity and disease. Nat Rev Neurosci 2013; 14(6): 383–400.

Papale LA, Beyer B, Jones JM, Sharkey LM, Tufik S, Epstein M, et al. Heterozygous mutations of the voltage-gated sodium channel SCN8A are associated with spike-wave discharges and absence epilepsy in mice. Hum Mol Genet 2009; 18(9): 1633–41.

Petrovski S, Kwan P. Unraveling the genetics of common epilepsies: approaches, platforms, and caveats. Epilepsy Behav 2013; 26(3): 229–33.

Pierson TM, Yuan H, Marsh ED, Fuentes-Fajardo K, Adams DR, Markello T, et al. GRIN2A mutation and early-onset epileptic encephalopathy: personalized therapy with memantine. Ann Clin Transl Neurol 2014; 1(3): 190–8.

Pioro EP, Brooks BR, Cummings J, Schiffer R, Thisted RA, Wynn D, et al. Dextromethorphan plus ultra low-dose quinidine reduces pseudobulbar affect. Ann Neurol 2010; 68(5): 693–702.

Rossi M, Chatron N, Labalme A, Ville D, Carneiro M, Edery P, et al. Novel homozygous missense variant of GRIN1 in two sibs with intellectual disability and autistic features without epilepsy. Eur J Hum Genet 2017; 25(3): 376–80.

Shahaf G, Marom S. Learning in networks of cortical neurons. J Neurosci 2001; 21(22): 8782–8.

Slopien A, Dmitrzak-Weglarz M, Rybakowski F, Rajewski A, Hauser J. [Genetic background of ADHD: genes of the serotonergic system, other candidate genes, endophenotype]. Psychiatr Pol 2006; 40(1): 33–42.

Traynelis SF. Software-based correction of single compartment series resistance errors. Journal of neuroscience methods 1998; 86(1): 25–34.

Traynelis SF, Wollmuth LP, McBain CJ, Menniti FS, Vance KM, Ogden KK, et al. Glutamate receptor ion channels: structure, regulation, and function. Pharmacol Rev 2010; 62(3): 405–96.

Vajda I, van Pelt J, Wolters P, Chiappalone M, Martinoia S, van Someren E, et al. Low-frequency stimulation induces stable transitions in stereotypical activity in cortical networks. Biophys J 2008;94(12): 5028–39.

Van Pelt J, Corner MA, Wolters PS, Rutten WL, Ramakers GJ. Longterm stability and developmental changes in spontaneous network burst firing patterns in dissociated rat cerebral cortex cell cultures on multielectrode arrays. Neurosci Lett 2004; 361(1-3): 86–9.

Wagenaar DA, Pine J, Potter SM. Effective parameters for stimulation of dissociated cultures using multielectrode arrays. J Neurosci Methods 2004; 138(1-2): 27–37.

Wagenaar DA, Pine J, Potter SM. An extremely rich repertoire of bursting patterns during the development of cortical cultures. BMC Neurosci 2006; 7: 11.

Wagenaar DA, Potter SM. A versatile all-channel stimulator for electrode arrays, with real-time control. J Neural Eng 2004; 1(1): 39–45.

Wagnon JL, Mahaffey CL, Sun W, Yang Y, Chao HT, Frankel WN. Etiology of a genetically complex seizure disorder in Celf4 mutant mice. Genes Brain Behav 2011; 10(7): 765–77.

Wyllie DJ, Behe P, Colquhoun D. Single-channel activations and concentration jumps: comparison of recombinant NR1a/NR2A and NR1a/NR2D NMDA receptors. J Physiol 1998; 510 (Pt 1): 1–18.

XiangWei W, Jiang Y, Yuan H. De Novo Mutations and Rare Variants Occurring in NMDA Receptors. Curr Opin Physiol 2018; 2: 27–35.

Yang M, Bozdagi O, Scattoni ML, Wohr M, Roullet FI, Katz AM, et al. Reduced excitatory neurotransmission and mild autism-relevant phenotypes in adolescent Shank3 null mutant mice. J Neurosci 2012; 32(19): 6525–41.

Yuan H, Erreger K, Dravid SM, Traynelis SF. Conserved structural and functional control of N-methyl-D-aspartate receptor gating by transmembrane domain M3. J Biol Chem 2005; 280(33): 29708–16.

Yuan H, Hansen KB, Vance KM, Ogden KK, Traynelis SF. Control of NMDA receptor function by the NR2 subunit amino-terminal domain. J Neurosci 2009; 29(39): 12045–58.

Yuan H, Low CM, Moody OA, Jenkins A, Traynelis SF. Ionotropic GABA and Glutamate Receptor Mutations and Human Neurologic Diseases. Mol Pharmacol 2015; 88(1): 203–17.

