## Supplementary Figures and Tables for "Modeling and treating *GRIN2A* developmental and epileptic encephalopathy in mice"

Supplementary Figure 1. Genetic transmission of S644G allele and husbandry of *Grin2a*$^{S644G}$ mice. Mendelian distribution of +/+, S644G/+, and S644G/S644G genotypes in (a) B6NJ and (b) F1 or F2 hybrid background. Observation gathered from multiple litters (>50). (c) Pups born from B6NJ S644G/+ females show no milk spot, indicative of poor maternal care.

**a.** B6NJ

♀ +/+ × ♂ S644G/+

| ***Grin2a* genotype** | **Obtained %** | **Expected %** |
|---|---|---|
| +/+ | 51% | 50% |
| S644G/+ | 49% | 50% |

♀ S644G/+ × ♂ S644G/+

| ***Grin2a* genotype** | **Obtained %** | **Expected %** |
|---|---|---|
| +/+ | * | 25% |
| S644G/+ | * | 50% |
| S644G/S644G | * | 25% |

* Mice born but did not survive past PND0

**b.** FVB/N × B6NJ

♀ +/+ × ♂ S644G/+

| ***Grin2a* genotype** | **Obtained %** | **Expected %** |
|---|---|---|
| +/+ | 47% | 50% |
| S644G/+ | 53% | 50% |

♀ S644G/+ × ♂ S644G/+

| ***Grin2a* genotype** | **Obtained %** | **Expected %** |
|---|---|---|
| +/+ | 23% | 25% |
| S644G/+ | 50% | 50% |
| S644G/S644G | 27% | 25% |

**c.**

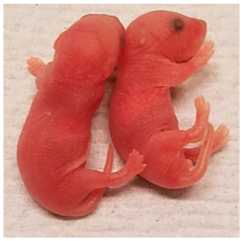

Supplementary Figure 2. Representative wake EEG of *Grin2a* S644G/+ and wildtype +/+ littermates. No obvious epileptiform activity was observed in traces from at least seven *Grin2a*$^{S644G/+}$ mice recorded for 48 hours each.

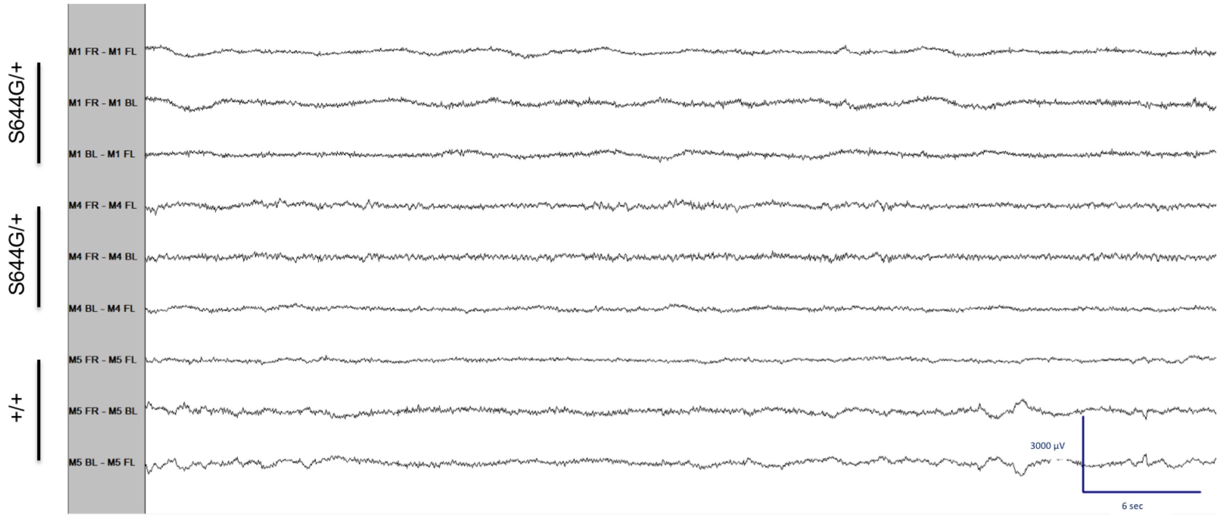

Supplementary Figure 3. GluN2A, GluN2B protein and mRNA in whole brain of two week old mouse pups. (a) Representative western blot of whole brain lysates probed for GluN2B, GluN2A, PSD95, β3-tubulin. (b) Plot of amount of protein normalized to β3-tubulin and wildtype. (c) mRNA expression quantification of total GRIN2B, GRIN2A, and PSD95 was determined by qRT-PCR and normalized to housekeeping gene, GADPH. Error bars indicate S.E.M. *p<0.05, **, ***, **** Student's |t|-test

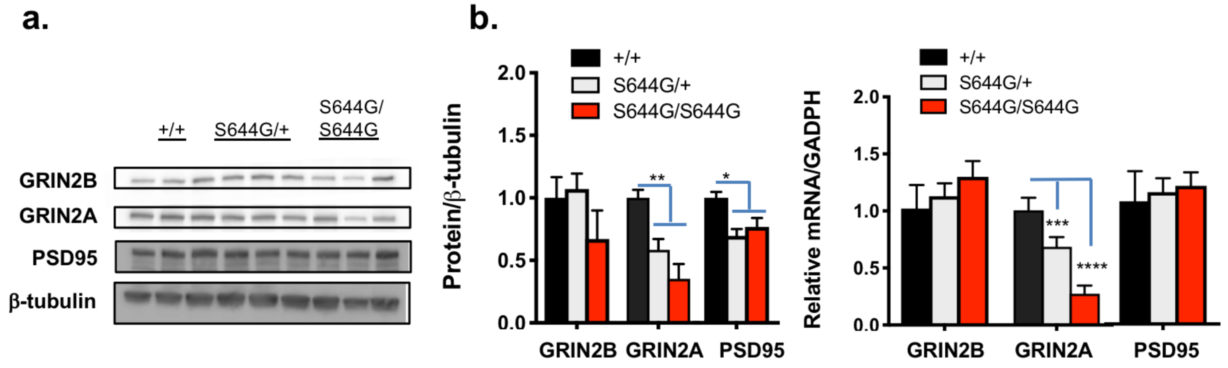

Supplementary Figure 4. Additional open field parameters – Ambulation (a) and center time (b) and vertical activity in B6NJ and $F_1$ hybrid mice comparing S644G/+ and +/+ (wildtype) genotypes. *p*-values shown are for the genotype effect in a 2-way repeated measures ANOVA.

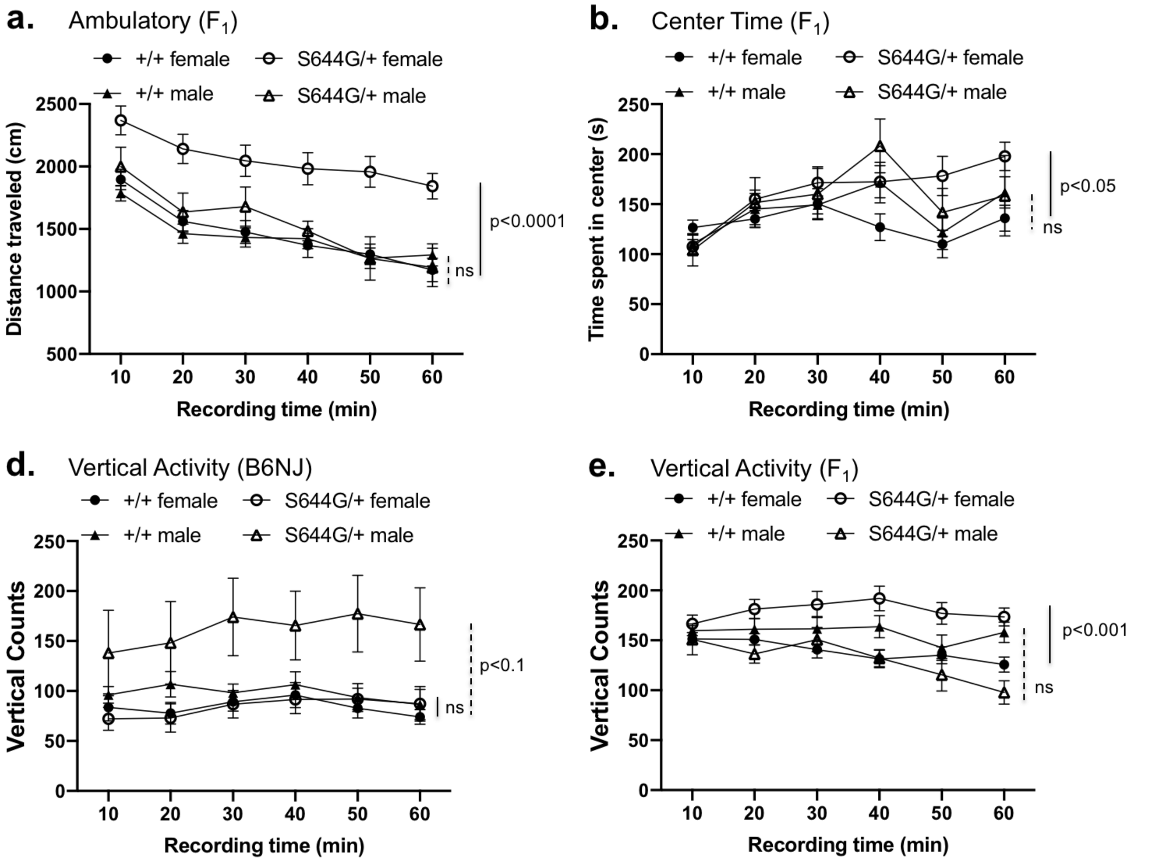

Supplementary Figure 5. Elevated plus maze. Elevated plus maze test was performed of S644G/+ and +/+ mice on both B6NJ and $F_1$ hybrid strain backgrounds, both sexes combined, showing the # of open arm entries (a), the percent time spent on the open arms (b), vs. total entries and time spent at the junction of the arms. There was a trend for S644G/+ mice to spend more time on the open arms (p=0.051, Mann-Whitney rank-sum test, strain backgrounds combined).

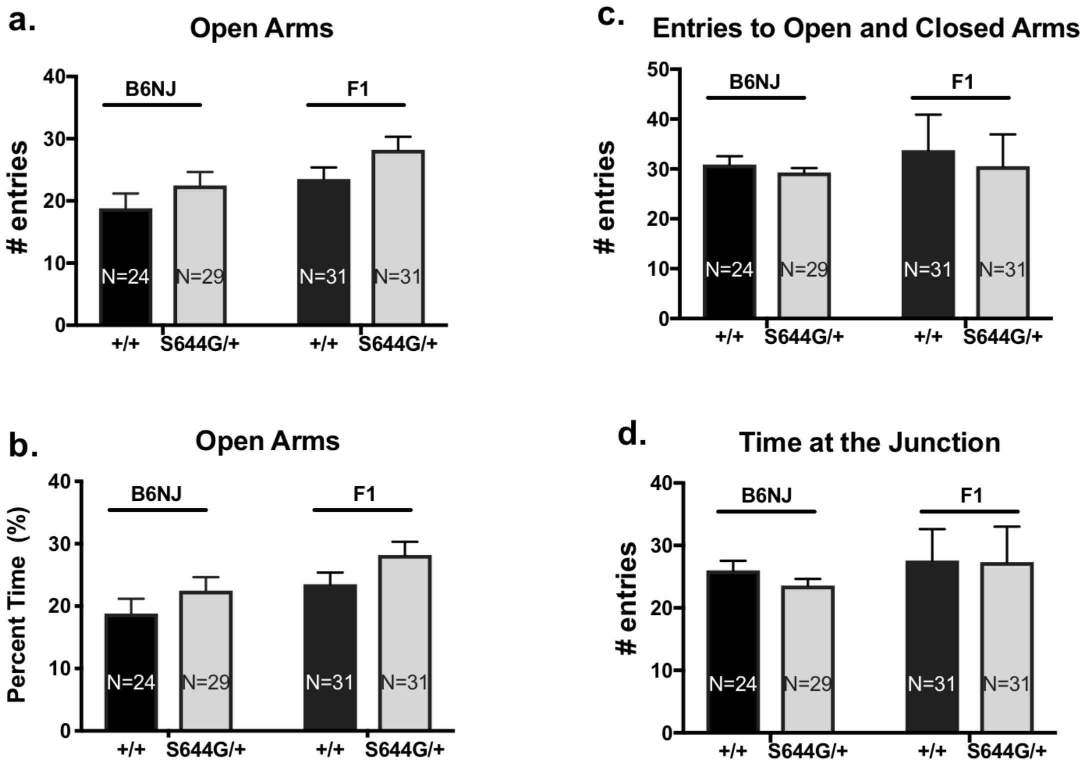

Supplementary Figure 6. a. Acoustic startle response ($F_1$ hybrid), b. representative ABR traces

As in the B6NJ line, heterozygous mice on the F1 background exhibited a markedly decreased response to a range of acoustic startle stimuli, without displaying impairments in the ABR test, indicating that reactivity to acoustic stimuli was altered, independent of defects in sensory perception.

**a.**

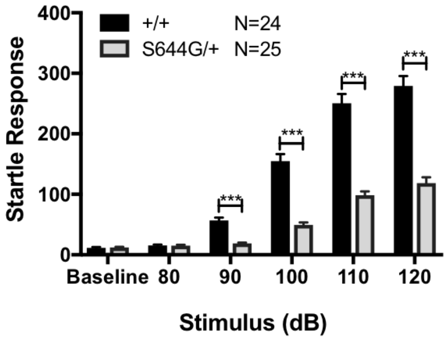


**b.**

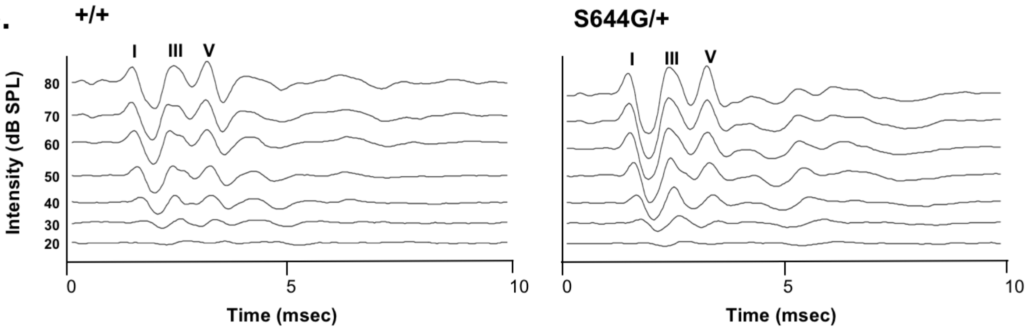

Supplementary Figure 7. a, b. Social interaction (B6NJ and $F_1$ hybrid), c, d. repetitive behaviors ($F_1$ hybrid)

As in the B6NJ line, heterozygous mice on the F1 background exhibited normal social interaction behaviors and significantly increase repetitive behaviors, especially self-grooming.

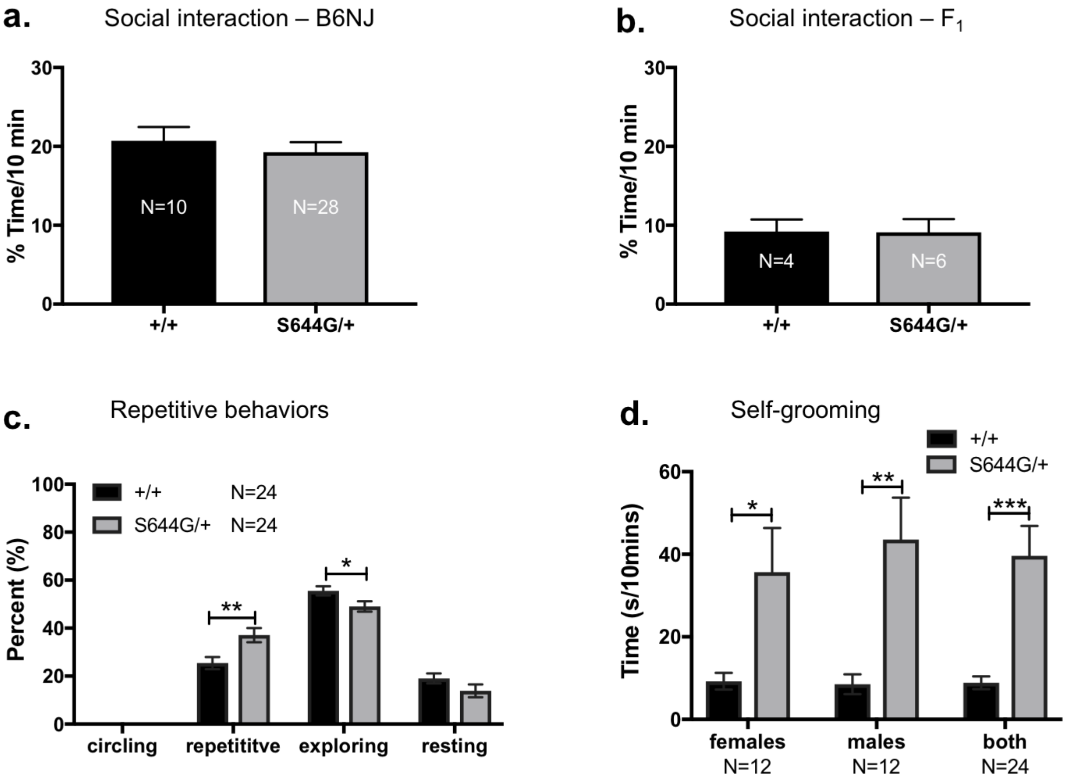

Supplementary Figure 8. Response of S644G-containing NMDARs to agonists *in vitro.* (a, b) Fitted composite glutamate and glycine concentration-response curves for diheteromeric human wildtype GluN1/GluN2A- and GluN1/GluN2A-S644G expressed in *Xenopus laevis* oocytes. Two-electrode voltage-clamp recordings were conducted at a holding potential of −40 mV. Glutamate potency was quantified as the half-maximally effective response ($EC_{50}$) of recombinant NMDARs receptors comprised of wild type or mutant GluNR2A in the presence of a maximally effective concentration of glycine (100 μM).Glycine potency was evaluated in the presence of a maximally effective concentration of glutamate (100 μM). (c, d) Fitted concentration-response curves for endogenous antagonists, $Mg^{2+}$ (at -60 mV) and $Zn^{2+}$(at -20 mV) at diheteromeric human GluN1/GluN2A NMDARs. (e,f) Fitted concentration-response curves for FDA-approved drugs (dextromethorphan, memantine) at diheteromeric human NMDARs. (g) Percentage current response at pH 6.8 versus 7.6 shows decreased current attenuation in the presence of increased proton concentrations (ie, low pH). Concentration-response curves were generated for co-agonists glutamate and glycine, channel blocker $Mg^{2+}$ and endogenous antagonist $Zn^{2+}$ to evaluate whether the mutation changes agonist potency or the sensitivity to endogenous modulators. (h) Agonist potency shifts by GluN2A-S644G-containing NMDARs *in vitro*. Fitted concentration-response curves for glutamate, in the presence of 100 μM glycine (a), and for glycine in the presence of 100 μM glutamate (i) for NMDARs that contain 0, 1, or 2 copies of the rat GluN2A-S644G variant expressed in *Xenopus laevis* oocytes. Results are mean $\pm$ SEM from 12-17 (glutamate) or 13-14 oocytes (glycine) from two separate injections. Either 1 or 2 copies of theGluN2A-S644G mutation significantly increases both glutamate and glycine potency of the NMDA receptors ($p<0.05$, ANOVA, Neuman-Kuels multiple comparison test).

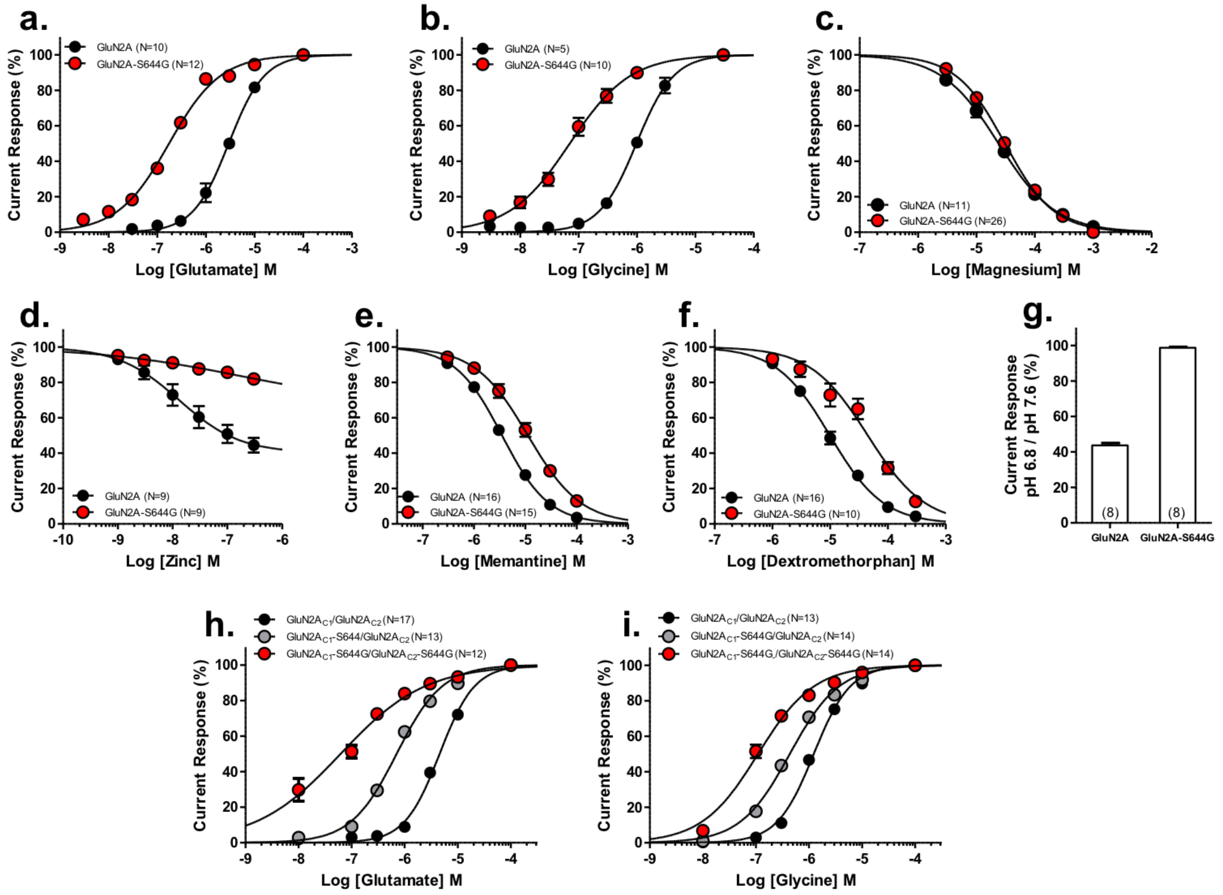
a.
GluN2A (N=10)
GluN2A-S644G (N=12)
Current Response (%)
Log [Glutamate] M
b.
GluN2A (N=5)
GluN2A-S644G (N=10)
Current Response (%)
Log [Glycine] M
c.
GluN2A (N=11)
GluN2A-S644G (N=26)
Current Response (%)
Log [Magnesium] M
d.
GluN2A (N=9)
GluN2A-S644G (N=9)
Current Response (%)
Log [Zinc] M
e.
GluN2A (N=16)
GluN2A-S644G (N=15)
Current Response (%)
Log [Memantine] M
f.
GluN2A (N=16)
GluN2A-S644G (N=10)
Current Response (%)
Log [Dextromethorphan] M
g.
Current Response pH 6.8 / pH 7.6 (%)
(8)
(8)
GluN2A
GluN2A-S644G
h.
GluN2A$_{C1}$/GluN2A$_{C2}$ (N=17)
GluN2A$_{C1}$-S644/GluN2A$_{C2}$ (N=13)
GluN2A$_{C1}$-S644G/GluN2A$_{C2}$-S644G (N=12)
Current Response (%)
Log [Glutamate] M
i.
GluN2A$_{C1}$/GluN2A$_{C2}$ (N=13)
GluN2A$_{C1}$-S644G/GluN2A$_{C2}$ (N=14)
GluN2A$_{C1}$-S644G,/GluN2A$_{C2}$-S644G (N=14)
Current Response (%)
Log [Glycine] M

Supplementary Figure 9. Additional multielectrode array analyses
(a) Temporal development of number of active electrodes, defined as electrodes recording at least 5 spikes per minute, demonstrates >94% active electrodes for each genotype by DIV13. (b) Mutant networks displayed significantly longer network bursts (NB) relative to wildtype networks. Error bars in (a) and (b) indicate SEM – statistical tests were done as described in Figure 6 and results listed on Supplementary Table 4. (c) Concentration-response curve of NMDA demonstrates sensitivity of mutant networks to NMDA-mediated excitation. Data for each well are normalized to the baseline firing recorded prior to agonist addition. Data were derived from 4 independent cultures with ≥ 17 wells for each genotype. (d) Peristimulus time histograms (400 ms) (see Materials and Methods) pre- and post-exposure to specified NMDA concentrations (n=6 wells). At baseline, mutant networks display prolonged evoked burst duration compared to wildtype with a mean AUC of 450 ± 10.7 compared to 330 ± 4.9 for wildtype. NMDA addition decreased evoked burst duration for both wildtype and mutant neurons in a concentration-dependent fashion. 1 µM NMDA; wildtype AUC 300 ± 5.4, S644G/S644G AUC 340 ± 8.8. 10 µM NMDA; wildtype AUC 250 ± 6.2 and S644G/S644G 140 ± 4.7.

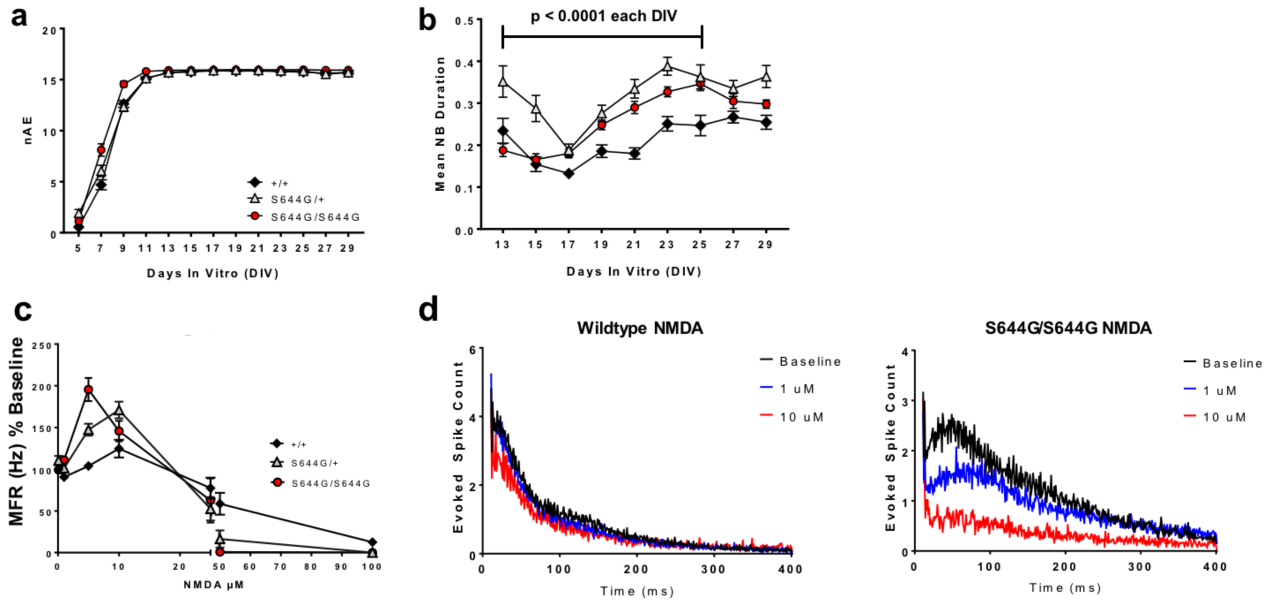

Supplementary Figure 10. Pharmacotherapy in the patient with GRIN2A S644G. Seizure frequency is shown during experimental treatment paradigm during which time the antiepileptic medications for the patient were altered.

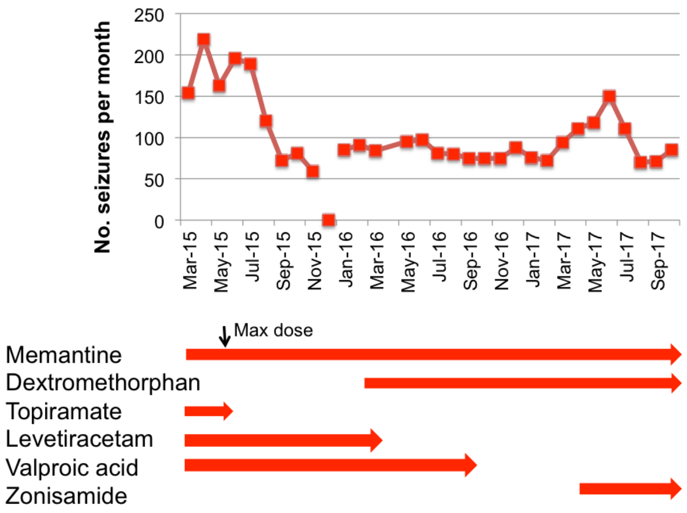

Supplementary Table 1. Agonist $EC_{50}$ and inhibitor $IC_{50}$ values for 0,1,or 2 rGluN2A-S644G-containing NMDARs in HEK cells.

| **Assay** | **GluN2A$_{C1}$/GluN2A$_{C2}$** | **GluN2A$_{C1}$-S644G/GluN2A$_{C2}$** | **GluN2A$_{C1}$-S644G/GluN2A$_{C2}$-S644G** |
|---|---|---|---|
| **Glu $EC_{50}$** | 4.5 $\mu$M (17; 4.0, 5.1) | 0.71 $\mu$M (13; 0.58, 0.88)* | 0.046 $\mu$M (12; 0.017, 0.12)* |
| **Gly $EC_{50}$** | 1.2 $\mu$M (13; 1.0,1.5) | 0.44 $\mu$M (14; 0.32, 0.61)* | 0.11$\mu$M (14; 0.0081, 0.15)* |
| **Dextromethorphan $IC_{50}$** | 13 $\mu$M (10; 11, 16) | 25 $\mu$M (11; 20, 29)* | 22 $\mu$M (13; 19, 25)* |
| **Memantine $IC_{50}$** | 4.4 $\mu$M (20; 3.0, 6.6) | 24 $\mu$M (10; 18, 34)* | 30 $\mu$M (11; 20, 44)* |

The data describing macroscopic currents from HEK cells transfected with the indicated cDNAs are expressed as mean (n; 95% CI determined from log $EC_{50}$ and $IC_{50}$ values). *, non overlapping confidence intervals.

Supplementary Table 2: Properties of NMDAR-mediated component of evoked EPSCs at the Schaffer collateral-CA1 pyramidal cell synapse

| | +/+ (n=15) | +/S644G (n=9) |
|---|---|---|
| Amplitude (peak, pA/pF) | 1.39 ± 0.25 | 1.56 ± 0.28 |
| 10-90% Rise time (ms) | 7.2 ± 0.4 | 7.6 ± 0.6 |
| $t_{FAST}$ (ms) | 50 ± 5.7 | 57 ± 9.2 |
| $t_{SLOW}$ (ms) | 306 ± 35 | 239 ± 23 |
| $t_{FAST}$ amplitude (%) | 74 ± 3.3 | 47 ± 5.1 |
| $t_W$ (ms) | 104 ± 4.1 | 150 ± 9.6 * |
| Charge transfer (pA ms/pF) | 163 ± 36 | 241 ± 36 |

Data are mean ±SEM. * indicates $p<0.05$ different by unpaired t-test for tau weighted. Power to detect an effect size of 1.5 (e.g. 30 ms change) was 0.92 (GPower 3.0).

Supplementary Table 3: Response time course of triheteromeric NMDARs containing 0, 1, or 2 copies of GluN2A-S644G

| | WT / WT (n=14) | S644G / WT (n=8) | S644G / S644G (n=9) |
|---|---|---|---|
| Amplitude (peak, pA/pF) | 57 ± 21 | 22 ± 5 | 36 ± 16 |
| Rise time (ms) | 9.9 ± 2.0 | 9.0 ± 1.4 | 8.8 ± 1.5 |
| $t_{FAST}$ deactivation (ms) | 55 ± 4.6 | 231 ± 38 | 547 ± 156 |
| $t_{SLOW}$ deactivation (ms) | -- | 1040 ± 253 | 2130 ± 415 |
| $t_{FAST}$ amplitude (%) | 100 | 72 | 66 |
| $t_W$ (ms) | 55 ± 4.6 | 456 ± 84* | 1220 ± 264*, $^{\#}$ |
| Charge transfer (pA ms/pF) | 2550 ± 691 | 10250 ± 3120* | 31000 ± 13100* |

Data are mean ± SEM. -- indicates not detected
WT/WT indicates HEK-293 cells transfected with GluN1 plus wildtype GluN2A$_{C1}$ and GluN2A$_{C2}$, S644G/WT indicates the same with 1 copy of GluN2A$_{C1}$-S644G and GluN2A$_{C2}$, and S644G/S644G indicates GluN2A$_{C1}$-S644G and GluN2A$_{C2}$-S644G.
*indicates $p<0.05$ by one-way ANOVA for log (tau weighted, $\mathbf{t_W}$) and charge transfer compared to WT/WT, $^{\#}$ indicates $p<0.05$ by one-way ANOVA for $\mathbf{t_W}$ compared to S644G/WT, statistical analysis was validated by Tukey's multiple comparison test as post-hoc analysis; power to detect an effect size of 3 (e.g. a 2-fold change) was 0.99 for $\mathbf{t_W}$ and 1.5 for charge transfer was 0.98 (G*Power 3.0.10)

Supplementary Table 4. Inhibitor $EC_{50}$ values for 0 or 2 rGluN2A-S644G-containing NMDARs in HEK cells.

| Name | Class | GluN2A $EC_{50}$ (N, 95% CI) | GluN2A-S644G $EC_{50}$ (N, 95% CI) |
|---|---|---|---|
| **Memantine** | Alzheimer's Disease | 3.5 μM (16; 2.7, 4.5) | 10.7 μM (15; 7.1,16) * |
| **Dextromethorphan** | Antitussive | 9.8 μM (16; 7.4, 13.1) | 43 μM (17; 26.9, 68.7) * |
| **Dextrorphan** | Metabolite | 1.1 μM (15; 0.9, 1.4) | 3.0 μM (17; 1.2, 7.4) |
| **Amantadine** | Antiviral | 131 μM (10; 87,197) | 182 μM (12; 131, 253) |
| **Ketamine** | Anesthetic | 6.8 μM (15; 4.4, 10.5) | 118 μM (5; 87, 161) * |
| **TCN-201** | GluN2A Competitive Antagonist | 0.22 μM (7; 0.17, 0.29) | 7.2 μM (6; 5.2, 9.9) * |
| **EU-1794-2** | Negative Allosteric Modulator | 0.62 μM (15; 0.47, 0.70) | 3.3 μM (17; 1.2, 3.4) * |

The data describing macroscopic currents from HEK cells transfected with the indicated cDNAs are expressed as mean (n; 95% CI determined from log $EC_{50}$). * non-overlapping confidence intervals.

Supplementary Table 5. Statistical analyses of MEA activity phenotypes.

| | Firing Rate | | | | Bursts per minute | | | |
|---|---|---|---|---|---|---|---|---|
| | Plate effect | Genotype effect | | | Plate | Genotype | | |
| DIV | *p*-val | *p*-val | *p*-val adj.† | vs. wt^ | raw only | *p*-val | *p*-val adj.† | vs. wt^ |
| DIV9 | 9.29E-30 | 6.02E-02 | ns | | 3.34E-25 | 3.75E-06 | 4.12E-05 | hom |
| DIV11 | 3.86E-19 | 8.06E-05 | 8.86E-04 | hom het | 6.92E-20 | 1.26E-05 | 1.38E-04 | hom |
| DIV13 | 4.65E-19 | 1.98E-09 | 2.18E-08 | hom het | 7.10E-19 | 2.08E-11 | 2.29E-10 | hom het |
| DIV15 | 1.40E-20 | 3.83E-10 | 4.21E-09 | hom het | 4.89E-16 | 1.39E-12 | 1.53E-11 | hom het |
| DIV17 | 2.94E-27 | 3.63E-09 | 3.99E-08 | hom het | 8.88E-10 | 1.68E-11 | 1.85E-10 | hom het |
| DIV19 | 6.34E-11 | 7.33E-10 | 8.06E-09 | hom het | 1.10E-07 | 1.23E-08 | 1.35E-07 | hom het |
| DIV21 | 5.50E-24 | 1.75E-06 | 1.93E-05 | hom het | 6.57E-21 | 3.33E-11 | 3.66E-10 | hom het |
| DIV23 | 5.92E-15 | 6.63E-02 | ns | | 1.79E-07 | 9.88E-11 | 1.09E-09 | hom het |
| DIV25 | 1.93E-06 | 1.11E-01 | ns | | 4.20E-09 | 1.84E-07 | 2.02E-06 | hom het |
| DIV27 | 9.75E-08 | 1.41E-03 | 1.55E-02 | hom | 6.79E-10 | 4.74E-03 | ns | |
| DIV29 | 1.53E-02 | 4.43E-01 | ns | | 1.29E-04 | 7.23E-01 | ns | |

| | Mutual Information | | | | Network Burst Duration | | | |
|---|---|---|---|---|---|---|---|---|
| | Plate effect | Genotype effect | | | Plate | Genotype | | |
| DIV | *p*-val | *p*-val | *p*-val adj.† | vs. wt^ | raw only | *p*-val | *p*-val adj.† | vs. wt^ |
| DIV9 | 1.46E-17 | 1.78E-12 | 1.96E-11 | hom | 9.07E-26 | 9.74E-01 | ns | |
| DIV11 | 6.47E-14 | 2.89E-17 | 3.18E-16 | hom | 1.00E-23 | 1.54E-01 | ns | |
| DIV13 | 3.27E-09 | 2.60E-23 | 2.86E-22 | hom | 3.75E-10 | 1.31E-07 | 1.44E-06 | hom het |
| DIV15 | 5.20E-14 | 5.98E-11 | 6.58E-10 | hom | 7.31E-05 | 3.71E-09 | 4.08E-08 | hom het |
| DIV17 | 2.54E-06 | 5.09E-11 | 5.60E-10 | hom | 1.27E-06 | 6.33E-09 | 6.96E-08 | hom het |
| DIV19 | 2.02E-15 | 5.57E-05 | 6.12E-04 | hom | 1.62E-12 | 5.81E-10 | 6.39E-09 | hom het |
| DIV21 | 1.47E-08 | 4.50E-05 | 4.95E-04 | hom | 5.35E-06 | 1.38E-18 | 1.52E-17 | hom het |
| DIV23 | 1.99E-12 | 4.00E-05 | 4.40E-04 | hom | 4.29E-09 | 6.18E-13 | 6.80E-12 | hom het |
| DIV25 | 2.54E-08 | 5.01E-01 | ns | | 2.17E-13 | 5.34E-07 | 5.88E-06 | hom het |
| DIV27 | 4.51E-10 | 4.14E-02 | ns | | 3.89E-07 | 1.23E-03 | 1.35E-02 | hom het |
| DIV29 | 1.44E-09 | 7.93E-04 | 8.72E-03 | hom | 8.05E-08 | 4.15E-04 | 4.57E-03 | hom het |

**†Bonferroni adjustment: *p*-value x 11 (number of DIV's evaluated for each feature)**
**^Post-hoc Dunnet's test for hom vs wt or het vs wt, *p* <0.05**

Supplementary Table 6. Compound $IC_{50}$ values for wildtype, S644G/+, and S644G/S644G cortical neural networks on MEA.

| Compound | WT | S644G/+ | S644G/S644G |
|---|---|---|---|
| **Dextromethorphan $IC_{50}$** | 2.35 μM (26; 1.9, 2.8) | 3.00 μM (27; 2.5, 3.6) | 5.51 μM (12; 4.1, 6.6) |
| **Memantine $IC_{50}$** | 4.51 μM (29; 3.9, 5.2) | 3.36 μM (46; 2.9, 3.9) | 2.05 μM (4; 1.5, 2.8) |

The data are expressed as mean (n; 95% CI). Concentration response curves (Figure 7c, d) were constructed from a sigmoidal function of nonlinear regression from which the IC50 was determined. See Figure 7 for statistics.
